# Virus-like particle delivery enables orthogonal genome editing *in vitro* and *in vivo*

**DOI:** 10.64898/2026.02.10.705092

**Authors:** Soochul Shin, Bae-Geun Nam, Eunji Park, Min Gu Kim, Chang Sik Cho, Yong-Woo Kim, Sang-Yeon Lee, Jeong Hun Kim, Sangsu Bae

## Abstract

The advent of CRISPR–Cas systems has revolutionized multiple fields, including basic science, biotechnology, and medicine. Central to this versatility is the use of programmable guide RNAs (gRNAs), which enable flexible and specific gene targeting. Building on this principle, various CRISPR-associated tools have been developed, including Cas9, base editors, prime editors, and gene-regulation platforms. However, because most CRISPR modalities share a common gRNA platform, the simultaneous use of multiple tools is constrained by interactions among different gRNAs, limiting orthogonal genome editing. This study presents a method for orthogonal multiplexed gene editing by packaging distinct CRISPR effectors and their corresponding gRNAs into separate virus-like particles (VLPs), thereby virtually eliminating the gRNA crosstalk. Furthermore, we demonstrate that this VLP-based strategy enables orthogonal, multiplexed gene editing *in vivo* in mouse eyes and ears. This platform expands the CRISPR toolkit by enabling simultaneous, non-interfering genetic manipulations in both cultured cells and living organisms.

## Main

CRISPR–Cas systems have revolutionized biological research by enabling precise genetic manipulation and engineering. Initially developed for genome editing^1-6^, CRISPR technology is now used across diverse fields, including diagnostics^7,8^, gene editing therapy^9,10^, genetic screening^11,12^, cellular imaging^13^, and the production of transgenic organisms^14^. This versatility derives from the use of guide RNAs (gRNAs), which provide both flexibility and specificity, enabling Cas effectors to reach virtually any genomic target. Moreover, by leveraging this concept, a wide range of CRISPR-based tools has been developed, in addition to CRISPR–Cas9, which can induce double-strand breaks (DSBs) at target sites. Comparatively, base editors (BEs), including the cytosine base editor (CBE)^15^ and adenine base editor (ABE)^16^, enable targeted single-nucleotide conversions without inducing DSBs. Meanwhile, prime editing (PE) allows precise insertions, deletions, and substitutions^17^. CRISPR activation (CRISPRa)^18,19^ and CRISPR interference (CRISPRi)^20^ extend the system to transcriptional regulation without altering DNA sequences. Overall, these approaches have created a powerful genetic toolkit that is broadly used in both basic research and clinical applications.

One of the major advantages of CRISPR technology is the associated ability to target multiple sites simultaneously via gRNAs. However, the simultaneous use of numerous CRISPR-based tools remains a technical challenge. In particular, when using Cas proteins from the same species, interactions between gRNAs (*i*.*e*., gRNA crosstalk) may occur. Indeed, for *Streptococcus pyogenes* Cas9 (SpCas9), except for the PE system, which uses a specific PE guide RNA (pegRNA), most CRISPR-associated tools share the same RNA platform: a single-guide RNA (sgRNA), resulting in gRNA crosstalk among SpCas9-based tools. To bypass this, the use of orthogonal Cas proteins from different species has been proposed, each recognizing distinct gRNA scaffolds^21^. For example, SpCas9 can be used in combination with other type II species, such as *Streptococcus thermophilus* Cas9 (StCas9) and *Staphylococcus aureus* Cas9 (SaCas9), or with type V Cas12 effectors, in the absence of gRNA crosstalk. However, such approaches also face the following limitations: type II Cas9 orthologs often have more stringent protospacer adjacent motif (PAM) requirements and exhibit lower editing activity than SpCas9; type V Cas12 proteins are difficult to generate as nickase variants, which hinders the creation of BE or PE.

Recently, Komor and colleagues demonstrated a multiplexed orthogonal base editor, named MOBE^22^, in human cells that used an engineered gRNA containing each RNA aptamer to recruit the corresponding effector protein, either adenosine deaminase or cytidine deaminase. Although this approach significantly reduced gRNA crosstalk, the MOBE system fundamentally requires the use of an identical Cas9 variant, such as a Cas9 nickase with a D10A mutation (*i*.*e*., nCas9(D10A)) for both ABE and CBE. Therefore, the MOBE system cannot simultaneously use mixed formats of wild-type Cas9 and nCas9(D10A)-based tools (*e*.*g*., BEs), nCas9(H840A)-based tools (*e*.*g*., PE, snuABE), and dCas9(D10A/H840A)-based tools (*e*.*g*., CRISPRa, CRISPRi). Furthermore, the MOBE system necessitates more distinct aptamer systems as the number of modalities increases. Additionally, the MOBE system requires components such as a coat protein of the RNA bacteriophage MS2 and CRISPR complexes, which may limit further *in vivo* application. Collectively, an alternative strategy is still needed that enables orthogonality across diverse Cas variants and is compatible with *in vivo* applications.

Thus, this study presents a virus-like particle (VLP)-based solution that demonstrates orthogonal genome editing without gRNA crosstalk in both *in vitro* and *in vivo* settings. Indeed, by packaging each gRNA and the associated corresponding Cas effector within a separate VLP, we physically and spatially segregate different CRISPR ribonucleoproteins (RNPs) to virtually eliminate interactions between various Cas effectors, such as Cas9 nuclease, BE, PE, and CRISPRa/i, while preserving the specific activity of each tool. Furthermore, we demonstrate that VLPs deliver CRISPR RNPs more effectively *in vivo* to mouse eyes and ears. This method substantially expands the CRISPR toolkit by enabling multiplexed genetic engineering in both cultured cells and mice.

## Results

### Optimization of virus-like particles for efficient genome editing

We first aimed to produce VLPs encapsulating Cas9, ABE, and CBE. Thus, to produce VLPs containing Cas9/gRNA RNP complexes, we constructed four plasmids respectively encoding Gag–Pol (VLP structural components), Gag–SpCas9, gRNAs, and VSV-G (envelope protein) (**Supplementary Table 1**). A protease cleavage site was inserted between Gag and SpCas9 to enable proteolytic processing (**Fig. 1a**). In this experiment, five different gRNAs were designed to target sites in *RNF2, EMX1, CD244, HEK2*, and *AAVS1*, respectively. Subsequently, all plasmids were co-transfected into production cells (Gesicle Producer 293T), and two days after transfection, VLP version 4^23^, comprising Cas9 and each gRNA, named Cas9–VLPs, was purified through a 0.45 μm PVDF filter. The filtrate was subsequently concentrated using PEG-it Virus Precipitation Solution. We then delivered each Cas9–VLP into HEK293T cells, amplified each target site from genomic DNA (gDNA) by PCR (**Supplementary Table 2**), and subjected the amplicons to high-throughput sequencing (or targeted deep sequencing) to assess the insertion and deletion (indel) frequencies. The results showed that each Cas9–VLP exhibited high gene editing activity of over 90%, except for the CD244 target (approximately 20%), suggesting that this version of Cas9–VLP typically has a strong delivery ability *in vitro* (**Fig. 1b** and **Supplementary Fig. 1a**).

**Fig. 1.**
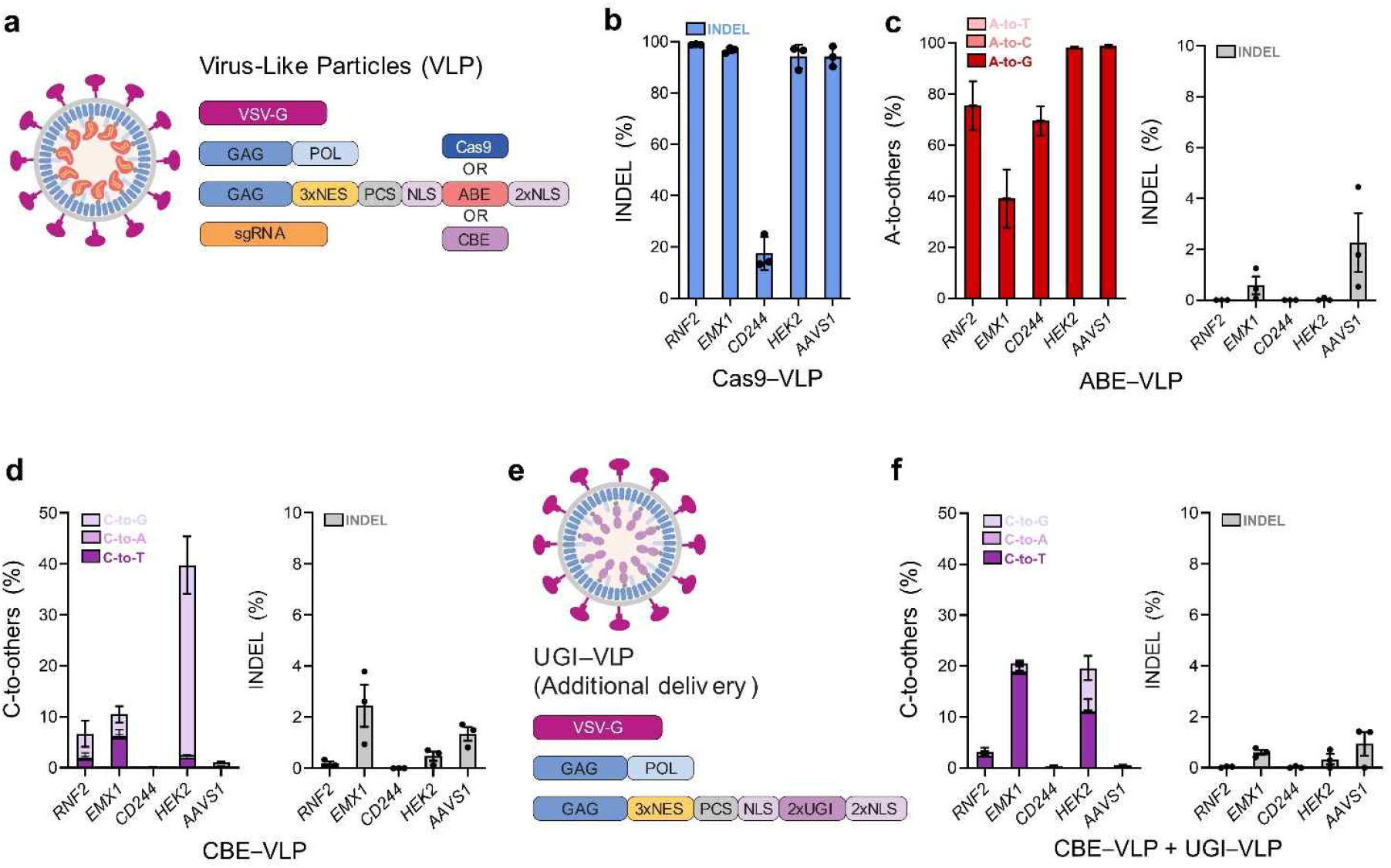
Optimization of virus-like particles for efficient genome editing. **a**, Schematic of the virus-like particle (VLP). **b**, Editing efficiencies for Cas9–VLP (15 μL) at the *RNF2, EMX1, CD244, HEK2*, and *AAVS1* loci in HEK293T cells. **c**, Editing efficiencies (left) and indel frequencies (right) for ABE– VLP (15 μL) at the *RNF2, EMX1, CD244, HEK2*, and *AAVS1* loci. **d**, Editing efficiencies (left) and indel frequencies (right) for CBE–VLP (15 μL) at the *RNF2, EMX1, CD244, HEK2*, and *AAVS1* loci. **e**, Schematic of UGI–VLP. Two copies of the uracil DNA glycosylase inhibitor (UGI) were fused to the C-terminus of the MMLV gag protein. **f**, Editing efficiencies (left) and indel frequencies (right) for CBE–VLP (15 μL) and UGI–VLP (25 μL) co-delivery at the *RNF2, EMX1, CD244, HEK2*, and *AAVS1* loci.

Similarly, we produced VLPs containing ABE8e and gRNA, termed ABE–VLPs, and treated HEK293T cells with each ABE–VLP designed to target the sites in *RNF2, EMX1, CD244, HEK2*, and *AAVS1*, respectively. Target sites were amplified by PCR, and A-to-others base conversion rates were evaluated through targeted deep sequencing. The results showed that ABE–VLPs exhibited dominant A-to-G base conversions with high activity exceeding 35% at all sites, suggesting that this version of the ABE–VLP also has reliable delivery capability *in vitro* (**Fig. 1c** and **Supplementary Fig. 1b**). Nonetheless, indels were also observed following the ABE–VLP treatment; however, the indel frequencies were less than 5% at all tested loci, markedly lower than the intended A-to-G conversions (**Fig. 1c**).

Finally, as with the ABE–VLPs, we produced VLPs containing AncBE4max and gRNA, termed CBE–VLPs, and treated HEK293T cells with each CBE–VLP designed to target sites in *RNF2, EMX1, CD244, HEK2*, and *AAVS1*, respectively. Target sites were amplified by PCR, and C-to-others base conversion rates were evaluated through targeted deep sequencing. However, the results showed that the CBE–VLPs exhibited overall low activity (<10%), and C-to-T base conversion was not the dominant pathway at most sites. For the *HEK2* site, C-to-others conversion rates were approximately 40%; however, C-to-T was only about 2% (**Fig. 1d** and **Supplementary Fig. 1c**). We hypothesized that the low editing efficiency and poor purity with the CBE–VLPs were due to the insufficient amount of the uracil DNA glycosylase inhibitor (UGI), a key component of CBE. UGI inactivates naturally occurring uracil DNA glycosylases (UDGs) in cells, thereby preventing uracil-containing DNA from being degraded and enhancing C-to-T editing activity. In our previous study, we observed that CBE RNPs exhibited reduced editing activity and low C-to-T purity, whereas supplementation with UGIs restored both editing activity and purity^25^. Therefore, using the same strategy, we additionally produced VLPs containing UGIs (**Fig. 1e**), termed UGI–VLPs, and delivered these UGI–VLPs alongside the CBE–VLPs; this resulted in improved C-to-T editing efficiency and purity. Across the five loci, C-to-T editing was detected at 2.3% in the *RNF2* target, 18.7% in the *EMX1* target, and 11.0% in the *HEK2* target, while C-to-T editing was low (approximately 0.2%) in both the *CD244* and *AAVS1* targets (**Fig. 1f**). Based on these results, the CBE–VLP was subsequently administered together with the UGI–VLP in future CBE applications.

### Implementation of orthogonal gene editing using virus-like particles

Given that the Cas9/gRNA RNP complex is tightly formed and rarely dissociates once formed^26-28^, we hypothesized that the gRNA crosstalk effect would not occur even if different VLPs were co-delivered. Notably, when multiple sets of Cas9/sgRNA1, ABE/sgRNA2, and CBE/sgRNA3 plasmids are simultaneously transfected into cells, gRNA cross-reactivity may be induced because these plasmids all share the same sgRNA platform, resulting in promiscuous mutations at each target (**Fig. 2a**). Conversely, when multiple VLPs of the Cas9–VLP with sgRNA1, ABE–VLP with sgRNA2, and CBE– VLP with sgRNA3 are simultaneously delivered into cells, we expect to enable orthogonal genome editing by inducing the intended editing events at each target site (**Fig. 2b**).

**Fig. 2.**
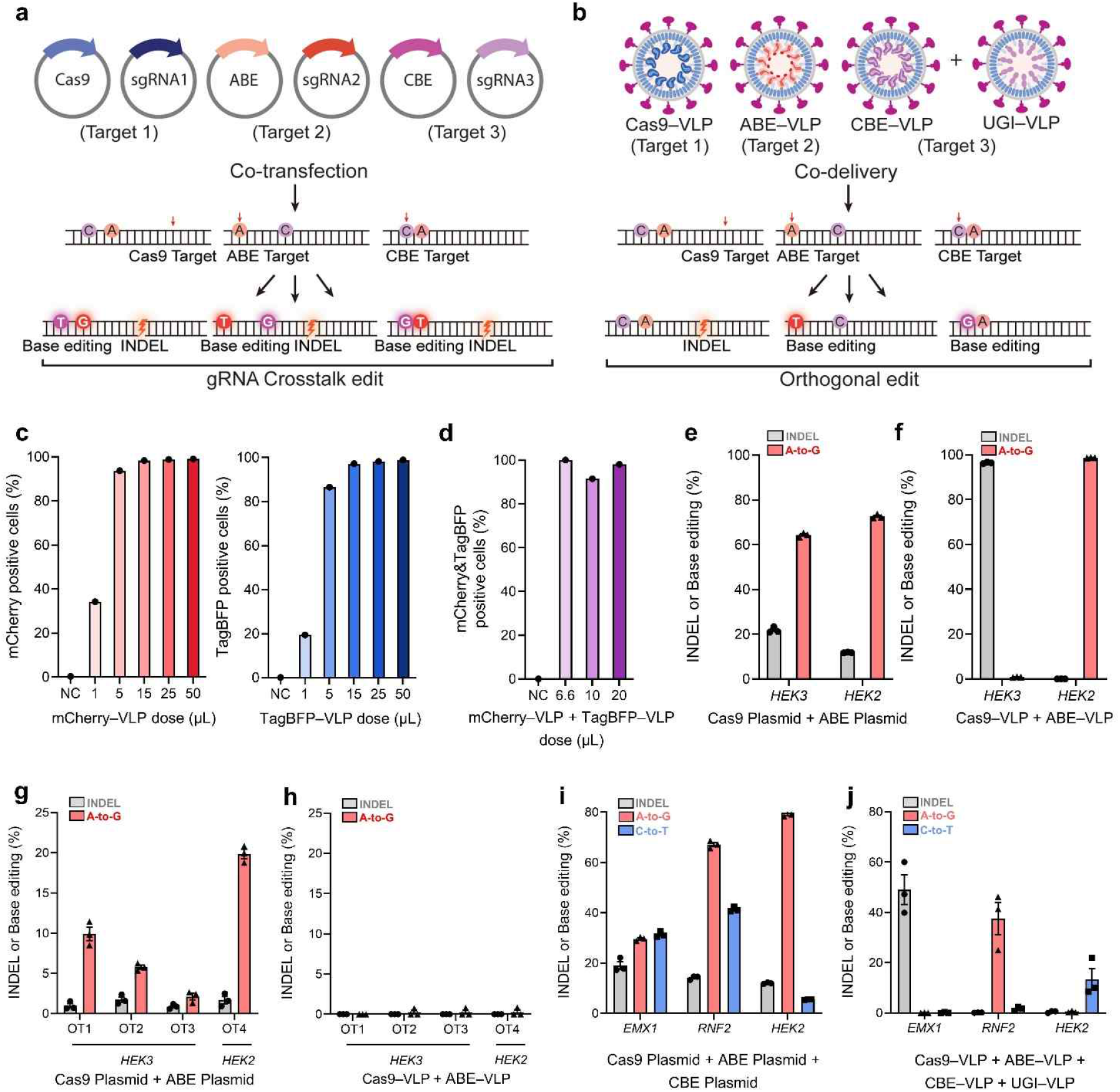
Implementation of orthogonal gene editing using virus-like particles. **a**, Overview of crosstalk during co-transfection of Cas9, ABE, and CBE plasmids with sgRNAs targeting their corresponding protospacers. **b**, Overview of orthogonal editing by co-delivery of Cas9–VLP, ABE– VLP, and CBE–VLP targeting their corresponding protospacers. **c**, Flow cytometry results of independent delivery with mCherry–VLP and TagBFP–VLP in HEK293T cells. **d**, Flow cytometry results of co-delivery with mCherry–VLP and TagBFP–VLP in HEK293T cells. **e**, Editing efficiencies following co-transfection of Cas9 and ABE plasmids with sgRNAs at the *HEK3* and *HEK2* loci. **f**, Editing efficiencies following co-delivery of Cas9–VLP (15 μL) and ABE–VLP (15 μL) at the *HEK3* and *HEK2* loci. **g**, Off-target frequencies at known *HEK3* and *HEK2* off-target sites after plasmid co-transfection. **h**, Off-target frequencies at known *HEK3* and *HEK2* off-target sites after VLP co-delivery. **i**, Editing efficiencies after the co-transfection of Cas9, ABE, and CBE plasmids at the *EMX1, RNF2*, and *HEK2* loci. **j**, Editing efficiencies after the co-delivery of Cas9–VLP (3 μL), ABE–VLP (7 μL), CBE–VLP (15 μL), and UGI–VLP (25 μL) at the *EMX1, RNF2*, and *HEK2* loci.

However, before conducting the orthogonal editing experiments, we investigated whether different types of VLPs could be delivered simultaneously and uniformly into cells. Thus, we performed a proof-of-concept study in which we generated VLPs loaded with red fluorescent protein (mCherry) or blue fluorescent protein (TagBFP), and delivered these either individually or in combination to HEK293T cells. Cells treated with increasing volumes (1, 5, 15, 25, and 50 µL) of mCherry–VLPs exhibited progressively stronger red fluorescence signals as measured by flow cytometry, with 34.2%, 93.7%, 98.3%, 98.8%, and 99.1%, respectively (**Fig. 2c**, left). Meanwhile, the TagBFP–VLPs exhibited a similar dose-dependent result, with the proportion of TagBFP-positive cells reaching 19.4%, 86.5%, 97.2%, 98.2%, and 98.9%, respectively (**Fig. 2c**, right). Interestingly, when both mCherry- and TagBFP-VLPs were delivered simultaneously with increasing volumes (total 6.6, 10, and 20 µL), a predominantly double-positive cell population was observed, demonstrating that cells could efficiently internalize multiplexed VLPs at once (**Fig. 2d** and **Supplementary Fig. 2**).

Next, we investigated whether VLP-mediated orthogonal editing occurs with two genome-editing tools: Cas9 and ABE. When two sets of plasmids—Cas9/sgRNA1 for *HEK3* target and ABE/sgRNA2 for *HEK2* target—were simultaneously transfected into HEK293T cells, both the indel and A-to-G conversion events were detected at both the *HEK3* and *HEK2* targets, indicating the occurrence of gRNA crosstalk between Cas9 and ABE (**Fig. 2e**). Conversely, when two types of VLPs —Cas9–VLPs containing sgRNA1 for *HEK3* target and ABE–VLPs containing sgRNA2 for *HEK2* target—were simultaneously delivered into HEK293T cells, exclusive indel formation was observed at the *HEK3* target alongside the intended A-to-G conversion at *HEK2* target (**Fig. 2f**), strongly suggesting the success of orthogonal editing between Cas9 and ABE. Notably, co-transfection with Cas9/sgRNA1 and ABE/sgRNA2 plasmids also led to orthogonal off-target editing. For example, A-to-G conversion was observed with up to 9.9% efficiency at the off-target sites of Cas9/sgRNA1, while indel formation was observed with up to 1.7% efficiency at the off-target site of ABE/sgRNA2 (**Fig. 2g**). In contrast, co-delivery of Cas9–VLP and ABE–VLP showed an overall very low off-target activity of less than 0.3% (**Fig. 2h**). As reported in previous studies^23,29^, the low off-target effect of RNP delivery was likely attributed to the associated relatively short duration and rapid degradation in cellular environments.

Subsequently, we extended the concept to three types of gene editing tools: Cas9, ABE, and CBE. Hence, we chose three target sites in *EMX1, RNF2*, and *HEK2*, where the Cas9–VLP, ABE–VLP, and CBE–VLP all exhibited activity, as validated in the above experiments (**Fig. 1**). When three sets of plasmids—Cas9/sgRNA1 for *EMX1* target, ABE/sgRNA2 for *RNF2* target, and CBE/sgRNA3 for *HEK2* target—were simultaneously transfected into HEK293T cells, the promiscuous indel, A-to-G, and C-to-T conversion events were detected at all targets, indicating extensive occurrence of gRNA crosstalk among Cas9, ABE, and CBE (**Fig. 2i**). However, when three types of VLPs—Cas9–VLP, ABE–VLP, and CBE–VLP to target *EMX1, RNF2*, and *HEK2* loci, respectively—were simultaneously delivered into HEK293T cells, the intended indel formation, A-to-G conversion, C-to-T conversion events were exclusively observed at the *EMX1, RNF2*, and *HEK2* targets, respectively (**Fig. 2j**), strongly supporting the success of orthogonal editing among Cas9, ABE, and CBE. Taken together, these data demonstrate multiplexed orthogonal genome editing via VLP-mediated RNP delivery of Cas9, ABE, and CBE.

### Extension of orthogonal editing to gene expression regulation

To further extend the usability of VLP-mediated orthogonal genome editing, we aimed to develop a PE system for VLP delivery—PE is a versatile tool that can induce all types of base conversions and indels within about 40 nucleotides. Therefore, in addition to Cas9 and BEs, the implementation of PE using VLPs is essential for expanding the diversity and scope of gene editing. Hence, we prepared PE2max and pegRNA constructs and encapsulated them in VLP version 3b (PE–VLP), which has previously been validated to increase internal cargo space and enhance the PE/pegRNA binding fraction^30^.

We produced ABE–VLPs to target the *HEK4* site and PE–VLPs to target the *HEK2* site (**Fig. 2a**), while also constructing two sets of plasmids—ABE/sgRNA1 for *HEK4* and PE/pegRNA1 for *HEK2*— as a control. When both the ABE/sgRNA1 and PE/pegRNA1 plasmids were co-transfected into HEK293T cells, A-to-G conversion events were detected at both the *HEK4* and *HEK2* target sites. In contrast, PE events were detected only at the *HEK2* target (**Fig. 3b**). This result suggests that gRNA crosstalk occurs only in one direction, from ABE to PE, because ABE can accommodate both sgRNA and pegRNA, leading to A-to-G editing at both the ABE and PE target sites. Alternatively, when ABE– VLP and PE–VLP were simultaneously delivered into HEK293T cells, exclusive orthogonal editing was observed: the A-to-G conversion frequency was 4.1% at *HEK4* with negligible PE efficiency, whereas the intended edit by PE was 1.9% at *HEK2* with undetectable ABE-mediated base conversion (**Fig. 3c**), indicating successful orthogonal editing between ABE and PE.

**Fig. 3.**
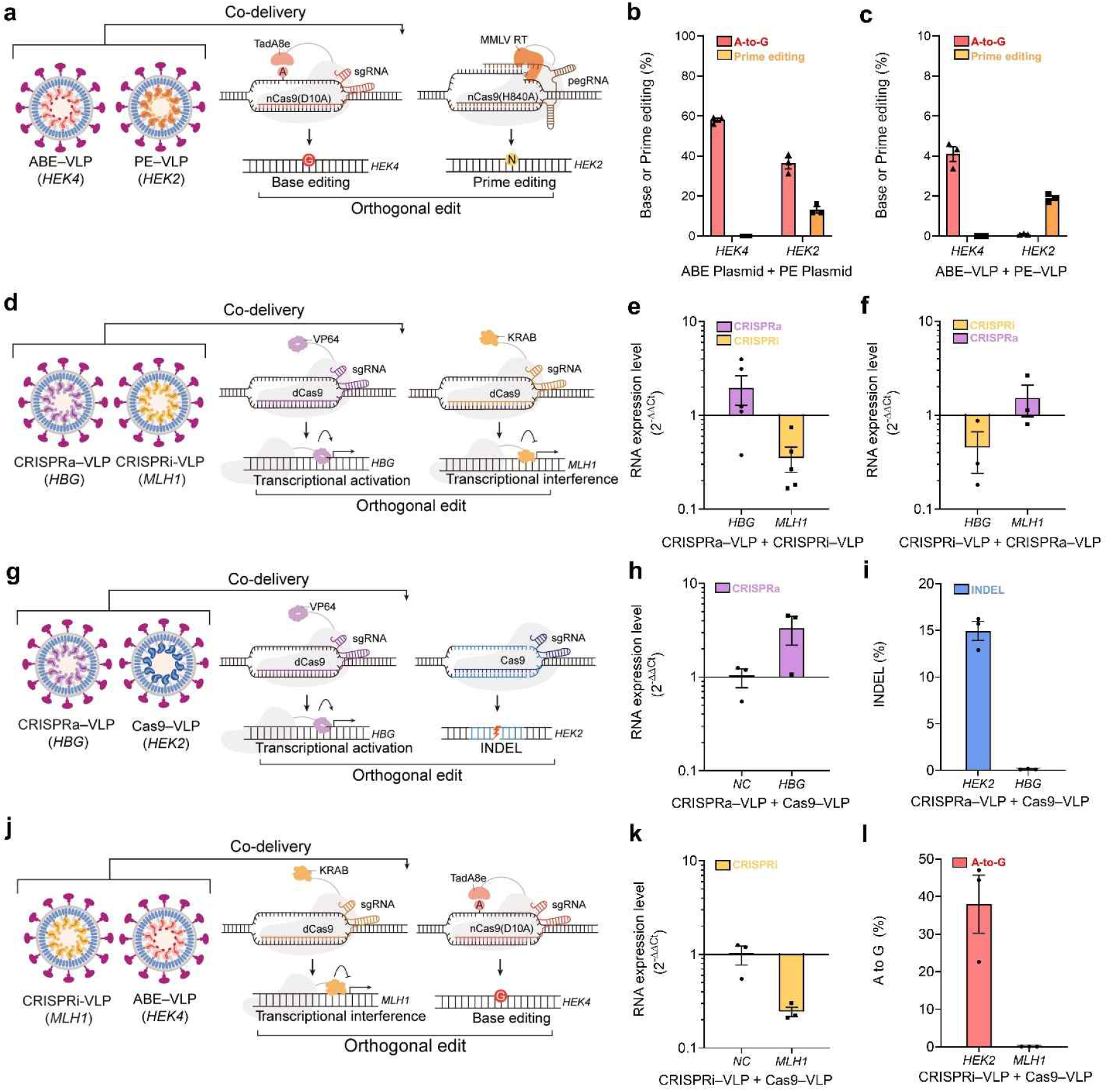
Extension of orthogonal editing to gene expression regulation. **a**, Overview of orthogonal editing of the ABE–VLP and PE–VLP at their corresponding protospacers. **b**, Editing efficiencies for the co-transfection of the ABE and PE plasmids with sgRNA and pegRNA at the *HEK4* and *HEK2* loci in HEK293T cells. **c**, Editing efficiencies for the co-delivery of ABE–VLP (15 μL) and PE–VLP (15 μL) at the *HEK4* and *HEK2* loci. **d**, Schematic of orthogonal targeting with CRISPRa–VLP and CRISPRi–VLP. **e**, mRNA expression changes in HeLa cells: upregulation of *HBG* by CRISPRa–VLP (30 μL) and downregulation of *MLH1* by CRISPRi–VLP (30 μL). Upon co-delivery, both effects were preserved independently, confirming orthogonality. **f**, Reversed targeting in which CRISPRi–VLP (30 μL) was directed to *HBG* and CRISPRa–VLP (30 μL) to *MLH1*: reduced *HBG* expression and increased *MLH1* expression. **g**, Schematic of orthogonal targeting of CRISPRa–VLP and Cas9–VLP. **h**, mRNA expression changes upon CRISPRa–VLP (*HBG*) (15 μL) and Cas9–VLP (*HEK2*) (5 μL) co-delivery at the *HBG* locus in HeLa cells. **i**, Indel frequencies of CRISPRa–VLP (15 μL) and Cas9–VLP (5 μL) co-delivery at the *HEK2* and *HBG* loci. **j**, Schematic of orthogonal targeting of CRISPRi–VLP and ABE– VLP. **k**, mRNA expression changes upon CRISPRi–VLP (*MLH1*) (15 μL) and ABE–VLP (*HEK2*) (5 μL) co-delivery at the *MLH1* locus in HeLa cells. **l**, Editing efficiencies of CRISPRi–VLP (15 μL) and ABE–VLP (5 μL) co-delivery at the *HEK2* and *MLH1* loci.

Next, we aimed to extend the VLP-based strategy for regulating gene expression. Thus, we established VLP version 4 encapsulating dCas9–VP64/sgRNA and dCas9–KRAB/sgRNA, named CRISPRa–VLP and CRISPRi–VLP, respectively (**Fig. 3d**). In this experiment, sgRNA1 was designed to target *HBG*, while sgRNA2 was designed to target *MLH1*. When two types of VLPs—CRISPRa-VLP for *HBG* and CRISPRi-VLP for *MLH1*—were delivered into HeLa cells, reverse transcription-quantitative PCR (RT-qPCR) analysis revealed that the expression level of *HBG* increased 2.0-fold compared to the control, while the expression level of *MLH1* decreased 0.4-fold compared to the control (**Fig. 3e**). Conversely, when we changed the target, *i*.*e*., CRISPRa–VLP for *MLH1* and CRISPRi–VLP for *HBG*, RT-qPCR analysis revealed that the expression level of *HBG* decreased 0.5-fold compared to the control, while the expression level of *MLH1* increased 1.5-fold compared to the control (**Fig. 3f**). These results strongly support the success of multiplexed orthogonal gene regulation between CRISPRa and CRISPRi.

Lastly, we examined whether gene editing and gene expression regulation could be induced in the absence of gRNA crosstalk effects. Hence, we first prepared CRISPRa–VLPs for *HBG* and Cas9–VLPs for *HEK2* (**Fig. 3g**). When CRISPRa–VLPs and Cas9–VLPs were simultaneously delivered into HeLa cells, we found that only the expression level of *HBG* was selectively increased by 3.3-fold compared to the control (**Fig. 3h**), and that indel formation was selectively observed only at the *HEK2* locus, with a frequency of 14.9% (**Fig. 3i**). Similarly, we next prepared CRISPRi–VLPs for *MLH1* and ABE–VLPs for *HEK2* (**Fig. 3j**). When CRISPRi–VLPs and ABE–VLPs were simultaneously delivered into HeLa cells, we observed a selective 0.2-fold decrease in *MLH1* expression compared with the control (**Fig. 3k**), and a 38.0% of A-to-G conversion that occurred selectively at the *HEK2* locus (**Fig. 3l**). Collectively, these results demonstrate the versatility of VLP-based orthogonal systems, highlighting their potential for efficient and independent multiplexed genetic engineering, including gene editing and gene regulation.

### Orthogonal genome editing using virus-like particles in mouse ears and eyes

*In vivo* genome editing represents a critical step toward therapeutic translation; however, additional barriers, including tissue accessibility and delivery efficiency, must still be overcome. Therefore, we investigated whether VLP-based delivery systems could maintain orthogonality beyond cultured cells, thereby assessing the feasibility of these systems as a versatile platform for *in vivo* therapeutic genome editing.

However, before conducting *in vivo* experiments, we first assessed whether VLP-based orthogonal editing strategies function efficiently in cultured mouse cells. To this end, Cas9–VLPs, ABE–VLPs, and CBE–VLPs were generated and tested in NIH3T3 cells. In this experiment, sgRNAs were designed to target five different loci in the mouse genome: *Vegfa, Rosa26, Dnmt1, Rpe65*, and *Ar*. Targeted deep sequencing results revealed that Cas9–VLP, ABE–VLP, and CBE–VLP all exhibited activity at the *Vegfa, Rosa26, Dnmt1*, and *Ar* loci (**Supplementary Fig. 3**). Moreover, we confirmed that orthogonal genome editing is robustly maintained in mouse cells upon co-delivery of Cas9–VLPs targeting *Vegfa* and ABE–VLPs targeting *Dnmt1* (**Supplementary Fig. 4**).

We then prepared Cas9–VLPs (2 x 10^9^ VP/μL) targeting *Vegfa* and ABE–VLPs (2 x 10^9^ VP/μL) targeting *Dnmt1*, and injected either or both into the inner ear of postnatal day 2 (P2) mice, using a total volume of 2 μL for each experiment. Cochlear tissues (*e*.*g*., organ of Corti) were collected from mice in the negative control (NC), single VLP-treated, and multiple VLP-treated groups on day 5 after injection (**Fig. 4a**). The indel frequency and A-to-G conversion frequency remained at baseline in the NC group: 0.2% and 0.5% at *Vegfa*, and 0.1% and 1.8% at *Dnmt1*, respectively. Meanwhile, in the single Cas9–VLP-treated group, the indel and A-to-G conversion frequencies at *Vegfa* were 5.0% and 0.7%, respectively, while in the single ABE-VLP-treated group, the frequencies at *Dnmt1* were 0.1% and 21.0%, respectively, (**Fig. 4b**). Conversely, in the combination-treated group (*i*.*e*., the group treated with both Cas9–VLP and ABE–VLP), the indel and A-to-G conversion frequencies at *Vegfa* were 3.2% and 0.6%, respectively, while the values at *Dnmt1* were 0.4% and 7.7%, respectively (**Fig. 4c**). These results strongly support the versatility and applicability of VLP-based orthogonal systems *in vivo* in mouse ears.

**Fig. 4.**
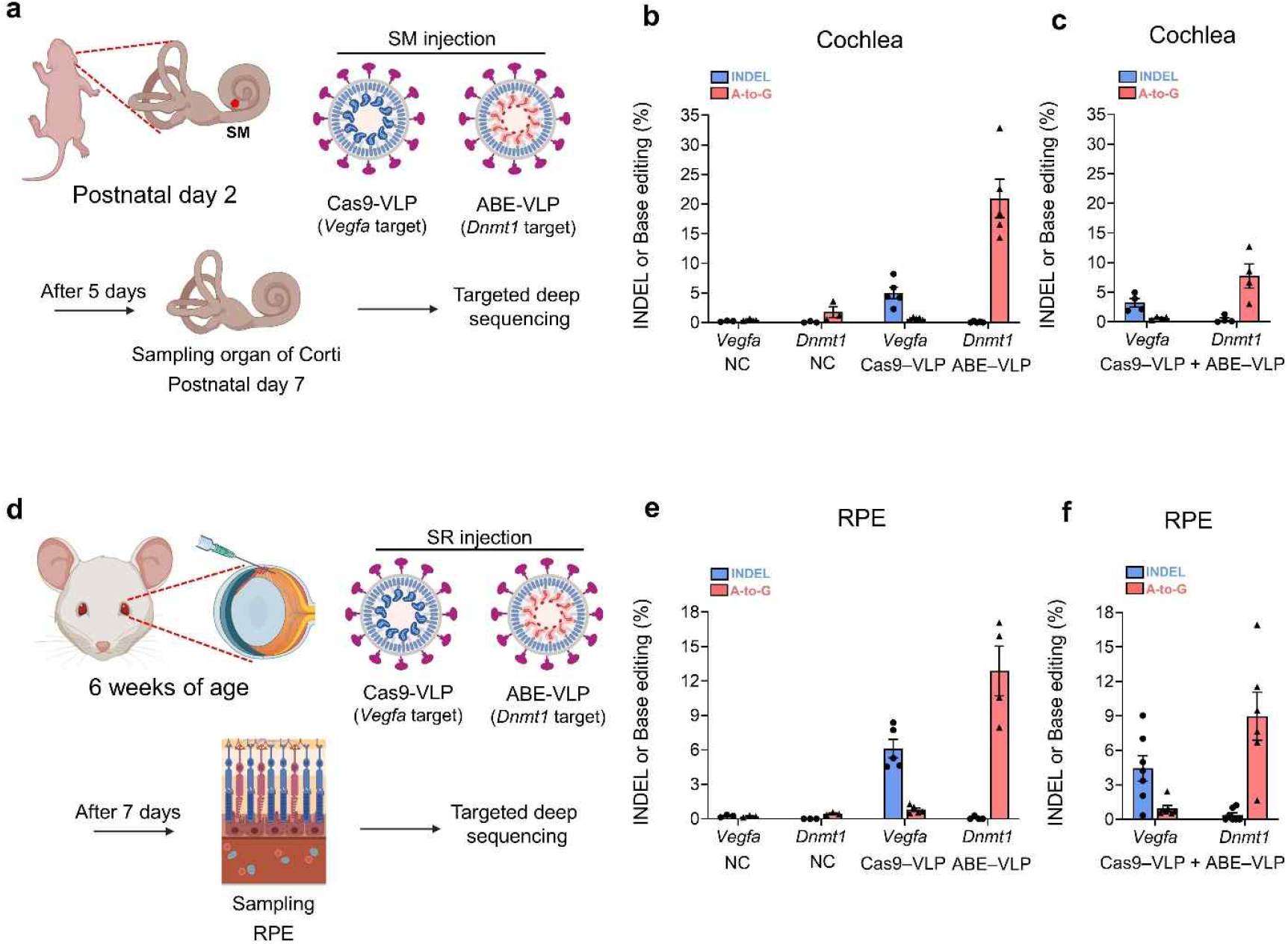
Orthogonal genome editing using virus-like particles in mouse ears and eyes. **a**, Schematic of cochlear scala media injection of a mixture of Cas9–VLP (*Vegfa*) and ABE–VLP (*Dnmt1*) in postnatal day 2 (P2) mice. Organ of Corti tissues were harvested at postnatal day 7 for sequencing analysis. **b**, Editing efficiencies at the *Vegfa* and *Dnmt1* loci after delivery of Cas9–VLP (2 μL) and ABE–VLP (2 μL) in the cochlea, respectively. **c**, Editing efficiencies at the *Vegfa* and *Dnmt1* loci after co-delivery of Cas9–VLP (1 μL) and ABE–VLP (1 μL) in the cochlea. **d**, Schematic of subretinal injection of Cas9– VLP (*Vegfa*) and ABE–VLP (*Dnmt1*) into 6-week-old mice, followed by sampling of the RPE for sequencing. **e**, Editing efficiencies at the *Vegfa* and *Dnmt1* loci after delivery of Cas9–VLP (2 μL) and ABE–VLP (2 μL) in the RPE, respectively. **f**, Editing efficiencies at the *Vegfa* and *Dnmt1* loci after co-delivery of Cas9–VLP (1 μL) and ABE–VLP (1 μL) in the RPE.

Next, we prepared Cas9–VLPs (2 x 10^9^ VP/μL) targeting *Vegfa* and ABE–VLPs (2 x 10^9^ VP/μL) targeting *Dnmt1*, and injected either or both into the subretinal region of the eyes of 6-week-old mice in a total volume of 2 μL for each experiment. Retinal pigment epithelium (RPE) was collected 7 days after injection to assess editing efficiency (**Fig. 4d**). In the NC group, the indel and A-to-G conversion frequencies remained at baseline: 0.3% and 0.2% at *Vegfa*, respectively, and 0.0% and 0.4% at *Dnmt1*, respectively. In the single Cas9–VLP-treated group, the indel and A-to-G conversion frequencies at *Vegfa* were 6.1% and 0.8%, respectively, whereas in the single ABE–VLP-treated group, the indel and A-to-G conversion frequencies at *Dnmt1* were 0.1% and 12.9%, respectively (**Fig. 4e**). Conversely, in the combination-treated group (*i*.*e*., the group treated with both Cas9–VLP and ABE–VLP), the indel and A-to-G conversion frequencies at *Vegfa* were 4.4% and 1.0%, respectively, while the values at *Dnmt1* were 0.4% and 9.0%, respectively (**Fig. 4f**). These results also support the applicability of the VLP-based orthogonal system in the mouse eye, demonstrating the versatility of *in vivo* gene editing in various tissue types.

## Discussion

This study demonstrates that the VLP-based delivery system enables multiplexed orthogonal genetic manipulation for sequence editing using Cas9, BE, and PE, and/or for gene-expression regulation using CRISPRa and CRISPRi. Our results reveal that packaging distinct CRISPR-associated RNP complexes into separate VLPs effectively prevents gRNA crosstalk, enabling simultaneous and independent manipulation of multiple genomic loci. This study also confirmed that multiplexed delivery of different CRISPR-associated tools can be achieved by simply adding more VLP types, without requiring additional system modifications or optimization steps. Moreover, the same VLP formulations optimized in *vitro* can be directly translated to *in vivo* applications, thereby offering a practical advantage for therapeutic development by reducing the gap between *in vitro* optimization and *in vivo* implementation.

Importantly, such orthogonal and multiplexed genome-editing capabilities are particularly relevant for the treatment of genetic disorders in which correction of multiple alleles or loci may be required. The prevalence of compound heterozygosity in autosomal recessive disorders is substantial; for example, approximately 40–50% of individuals with *GJB2*-related nonsyndromic hearing loss carry two distinct pathogenic alleles (52.9%, 55/104)^31^. Although correcting a single allele can often restore sufficient gene function, several exceptions exist. For example, in *ERCC2* (XPD)-associated *xeroderma pigmentosum*, each mutant allele disrupts separate helicase domains, so biallelic correction is required to restore the nucleotide-excision repair capacity fully^32^. Moreover, random monoallelic expression may render cells lacking biallelic correction functionally deficient^33^. In addition, this platform enables simultaneous gene editing and concurrent gene regulation, allowing combinatorial control of gene disruption, correction, activation, and suppression within the same biological system. Such an integrated strategy is particularly valuable for dissecting complex genetic networks and for designing therapeutic strategies that require coordinated modulation of multiple genes.

While our findings are promising, several areas warrant further investigation. First, although we improved the efficiency and purity of the CBE–VLPs by delivering additional UGI–VLPs, certain targets still exhibited suboptimal editing efficiency, indicating that full optimization of CBE–VLP performance has not yet been achieved. Additionally, the development of more sophisticated VLP formulations for quality control and methods to increase VLP titers is necessary to enhance the efficiency and applicability of this technology further. In the context of *in vivo* applications, organ- or tissue-specific improvements in delivery efficiency also remain critical. Furthermore, our study did not explore disease-relevant *in vitro* or *in vivo* models, which will be essential for assessing the therapeutic potential and translational feasibility of orthogonal VLP-mediated genome editing. Therefore, future studies should aim to address these aspects by applying this platform to clinically relevant disease models.

In conclusion, our study establishes VLPs as a versatile and effective platform for orthogonal genome editing in both cultured cells and mice. Indeed, by leveraging the natural properties of VLPs as delivery vehicles, we have developed a system that enables simultaneous, non-interfering application of multiple CRISPR modalities, thereby allowing more precise and scalable gene-editing applications in research and medicine.

## Acknowledgements

This work was supported by grants from the National Research Foundation of Korea (RS-2023-00211474 and RS-2024-00461016 to S.S.; RS-2024-00404132 to B.-G.N.; 2021M3A9H3015389, RS-2024-00451880, RS-2024-00455559, and SRC-NRF2022R1A5A102641311 to S.B.), grants from the Korean Fund for Regenerative Medicine (KFRM) (No. RS-2024-00332601), the Ministry of Food and Drug Safety (No. 25202MFDS003) in 2025, the SNUH Lee Kun-hee Child Cancer & Rare Disease Project (No. 25B-001-0700) to S.B., grants from the Phase III (Postdoctoral fellowship) of the SPST (SNU–SNUH Physician Scientist Training) Program and the Korea Health Technology R&D Project through the Korea Health Industry Development Institute (KHIDI) (No. RS-2025-02213987) to S.-Y.L., and grants from Seoul National University Hospital Research Grant (18-2023-0010) and the National Research Council of Science and Technology (24022-000) to J.H.K.

## Author contributions

S.S. and S.B. conceived the study; S.S., B.-G.N., and E.P. performed the cell experiments, and Y.-W.K. assisted; S.S., B.-G.N., and E.P. prepared VLPs, and M.G.K. and C.S.C. performed the in vivo mouse experiments under the supervision of S.-Y.L. and J.H.K.; S.B. supervised all procedures; S.S., B.-G.N., E.P., and S.B wrote the manuscript with assistance from other authors.

## Methods

### Cell culture conditions

HEK293T, HeLa, and NIH3T3 cells were cultured in Dulbecco’s Modified Eagle Medium (DMEM; Welgene, LM001-05) supplemented with 10% fetal bovine serum (FBS; Welgene, PK004) and 1% antibiotics (Welgene, LS203-1). Cells were maintained in a humidified incubator at 37 °C with 5% CO^2^. For cell passaging, cells were washed with Dulbecco’s phosphate-buffered saline (DPBS; Welgene, LB001-01) and detached using trypsin–EDTA (Welgene, LS015-01). After detachment, DMEM supplemented with 10% FBS was added to inactivate trypsin–EDTA, and the cell suspension was centrifuged at 100 × g for 3 min. The supernatant was aspirated, and the pellet was resuspended in fresh DMEM containing 10% FBS and 1% antibiotics, then transferred to a new culture dish to maintain proliferation.

### Plasmid construction

For Cas9–, ABE–, CBE–, CRISPRa–, and CRISPRi–VLP production, custom plasmids (pGag–SpCas9, pGag–ABE8e, pGag–AncBE4max, pGag–VP64–dCas9, and pGag–KRAB–dCas9) were constructed using pCMV_SpCas9 (Addgene #188489), pCMV_ABE8e (Addgene #138489), pCMV_AncBE4max (Addgene #112094), pCMV_VP64-dCas9 (Addgene #177171), and pCMV_KRAB-dCas9 (Addgene #110820), respectively, and these plasmids were also used in transfection experiments. Additionally, P4-PE2max (Addgene #211375) was used to produce PE–VLP. Custom or commercial plasmids were then transformed into 50 μL of DH5α competent cells. Single transformed colonies were inoculated into lysogeny broth (LB) medium containing antibiotics. Plasmids were isolated from cells using a DNA prep kit (Enzynomics, EP101-200N). A complete list of the custom plasmid sequences used in this study is provided in **Supplementary Table 1**.

### Transfection and genomic DNA extraction

The day before plasmid DNA transfection, HEK293T cells were washed with DPBS, detached with 0.05% trypsin–EDTA, and neutralized in DMEM containing 10% FBS and 1% antibiotics. Cells were then counted and seeded into a 48-well cell culture plate (SPL, 30048) at 3 × 10^4^ cells per well. SpCas9 expression plasmids (375 ng) and sgRNA expression plasmids (125 ng) were mixed with 50 μL of jetOPTIMUS (Polyplus-transfection, Illkirch, France) buffer and 0.5 μL JetOPTIMUS reagent and incubated at room temperature for 10 min. The prepared mixtures were added to the seeded 48-well plate. After 48 hours, the plate media was removed, and the transfected cells were washed with DPBS. The washed cells were lysed in 20 μL of lysis buffer (40 mM Tris (pH 8.0), 1% Tween-20, 0.2 mM EDTA, 0.2% Nonidet P-40, and 2 mg/mL of protease K) and incubated at 60 °C for 15 minutes for proteinase K activation, followed by 95 °C for 5 minutes for proteinase K inactivation.

### Production of VLPs

All v4 eVLPs were produced as previously described^23^. Briefly, Gesicle Producer-293T cells were plated at a density of 5 × 10^6^ cells per T75 flask (Corning, 353136) in 10 mL of DMEM with 10% FBS media and 1% antibiotics. After 20–24 hours, a mixture of plasmids was transfected into producer cells with jetPRIME transfection reagent (Polyplus, 101000001) following the manufacturer’s protocol. The plasmid composition for V4 eVLP was as follows: VSV-G (400 ng), wild-type MMLV Gag-Pol (3375 ng), Gag–SpCas9 (1125 ng), and sgRNA (4400 ng). For v3b eVLP, the plasmid composition included VSV-G (400 ng), wild-type MMLV Gag–Pol (2813 ng), Gag–COM–Pol (2000 ng), Gag–P3–Pol (422 ng), P4–SpCas9 (422 ng), and sgRNA (4400 ng). Then, 48 hours after transfection, supernatants were collected, centrifuged at 500 × g for 5 min, and filtered through a 0.45 μm polyvinylidene difluoride (PVDF) filter. For the eVLPs used in the *in vitro* experiments, 5X PEG-it Virus Precipitation Solution (System Biosciences, LV825A-1) was added to the supernatant and incubated overnight at 4 °C to precipitate the eVLPs. The next day, the eVLPs were pelleted by centrifugation at 1500 × g and 4 °C for 30 min and were concentrated 100-fold by resuspending in 100 μL of Opti-MEM (Life Technologies; 31985070). For the cell culture optimization experiments, all eVLPs were concentrated in the same manner to enable direct comparison of potency at equal volumes. In the *in vivo* experiments, eVLPs were concentrated by ultracentrifugation through a 20% (w/v) sucrose cushion in PBS at 141,000 × g and 4 °C for 2 hours, using an SW28 rotor in an Optima XPN Ultracentrifuge (Beckman Coulter). Following ultracentrifugation, the eVLP pellets were resuspended in cold Opti-MEM 100 μL per T75 flask.

### Calculating VLP titer

Measurement of MLV p30 within VLPs was performed using the MuLV Core Antigen ELISA kit (Cell Biolabs; VPK-156) according to the manufacturer’s protocols. The concentration of the VLP-associated p30 protein was calculated with the assumption that 20% of the observed p30 in solution was associated with VLPs, as was previously reported for MLV particles^34^. To estimate the number of SpCas9 protein molecules per VLP, we assumed that each VLP contained 1800 copies of p30, consistent with previous reports for MLV particles^34^.

### Targeted deep sequencing

Each target site was amplified in two sequential PCR rounds. In the first round (PCR1), target sites were amplified using primers containing Illumina forward and reverse sequencing adaptors. PCR1 was performed using 1 μL of genomic DNA extract and Sol Taq 2X Polymerase (Solg™ 2X Taq PCR Smart mix 1, w/o Band Doctor, STD01-M50h) under the following conditions: 95 °C for 2 min, followed by 24 cycles of 95 °C for 20 s, 56 °C for 40 s, and 72 °C for 30 s. The resulting PCR1 product was used as the template for a second amplification (PCR2) with Illumina index primers. PCR2 was performed using 1 μL of PCR1 product under the following cycling conditions: 95 °C for 2 min, followed by 29 cycles of 95 °C for 20 s, 56 °C for 40 s, and 72 °C for 30 s. Amplified products were verified by electrophoresis on a 1.5% agarose gel (Condalab, 8100). PCR2 products were pooled and purified with a PCR purification kit (GeneAll, 103-150). The libraries were sequenced on an Illumina MiniSeq platform, and the data were analyzed using Cas-Analyzer^35^, BE-Analyzer^36^, and PE-Analyzer^37^. A complete list of primers used in this study is provided in **Supplementary Table 2**.

### RNA isolation

Total RNA was extracted from cells transfected *in vitro* and *in vivo*, including cochlea and RPE samples, using TRIzol reagent (Invitrogen Life Technologies, Carlsbad, CA, U.S.A.). Isolated RNA samples were air-dried at room temperature for 30 min to allow remaining washing solution to evaporate, and resuspended in DEPC water. RNA quantification was performed using an Agilent 2100 Bioanalyzer (Agilent Technologies, Palo Alto, CA, U.S.A.) with the A260/A280 ratio.

### cDNA synthesis and RT-qPCR

For cDNA synthesis, 2000 ng of RNA was mixed with the PrimeScript™ RT reagent kit (TaKaRa, RR037B) and distilled water (DW) to a final volume of 20 μL. The reaction was conducted at 30 °C for 10 min, 42 °C for 60 min, 99 °C for 5 min, and 4 °C for 5 min. For qPCR, 10 μL of iTaq Universal SYBR Green Supermix (Bio-Rad, 1725124), 0.5 μL of each forward and reverse primer (20 pmol), and 9 μL of diluted cDNA were added to a 96-well qPCR plate. Then, qPCR was performed using the CFX96 Real-Time PCR Detection System (Bio-Rad) under the following cycling conditions: 95 °C for 3 min, followed by 65 cycles of 95 °C for 10 s, 55 °C for 10 s, and 72 °C for 30 s. A complete list of primers used in this study is provided in **Supplementary Table 2**, which also includes *GAPDH* as an internal control.

### Fluorescence-activated cell sorting (FACS)

For flow cytometry, cells were harvested, washed twice with cold PBS, and resuspended in FACS buffer (PBS supplemented with 2% FBS). CopGFP expression was detected using the FITC-A or Alexa Fluor 488 channel, mCherry expression was detected in the mCherry channel, and TagBFP expression was detected in the Pacific Blue channel. Cells were analyzed using a BD FACS Canto 8 flow cytometer (BD Biosciences) or FACSymphony A5 (BD Biosciences). All FACS data were processed using FlowJo software (TreeStar).

### Cochlear injection in mice and tissue collection

All animal procedures were approved by the Institutional Animal Care and Use Committee (IACUC) of Seoul National University College of Medicine. The humanized *MPZL2* mouse lines (*hMPZL2*^Q74X^ and *hMPZL2*^WT^; C57BL/6N background) were generated as described previously^38^. Mice of both sexes were randomly selected for inner-ear microinjections. Surgical procedures and injections were performed in accordance with published methods^38^, with minor modifications. Briefly, P2 *hMPZL2*^Q74X/WT^ and *hMPZL2*^WT/WT^ pups received VLP injections. Anesthesia was induced by hypothermia on ice, and a retroauricular incision was made under a surgical microscope (OPMI pico, ZEISS). Cas9–VLP (*mVegfa*, 2 μL) or ABE–VLP (*mDnmt1*, 2 μL) was injected separately (n = 5); a mixture of Cas9–VLP (*mVegfa*, 1 μL) and ABE–VLP (*mDnmt1*, 1 μL) was co-injected into the cochlear scala media (n = 4) using a Nanoliter 2020 Microinjection System (World Precision Instruments, Hertfordshire, UK). The total volume was 2 μL per cochlea, delivered at 7 nL/s using a MICRO2T SMARTouch microinjection controller (World Precision Instruments, Hertfordshire, UK). Incisions were closed with 6–0 nylon (NB617P, Ailee). Pups were warmed on a 37 °C heating pad for ∼5 min and returned to their mothers. For targeted deep sequencing, genomic DNA was purified from the organ of Corti of injected *hMPZL2*^Q74X/WT^ or *hMPZL2*^WT/WT^ mice at P7, as well as from untreated *hMPZL2*^WT/WT^ mice (negative control).

### RPE injection in mice and tissue collection

Six-week-old male C57BL/6 mice (Coatech, Gyeonggi-do, Korea) were used in this study. A total of six mice were allocated to each experimental group. Following anesthesia, ocular injections were performed subretinally (3 μL) to deliver PBS, Cas9–VLP (*mVegfa*), ABE–VLP (*mDnmt1*), and a mixture of Cas9–VLP (*mVegfa*) and ABE–VLP (*mDnmt1*). All drug formulations were prepared at a concentration of 2 × 10^9^ vp/μL. Seven days after injection, the eyes were enucleated, and the RPE was carefully dissected for further analysis. All animal procedures were conducted in accordance with the ARVO Statement for the Use of Animals in Ophthalmic and Vision Research.

## Data availability

All data are available from the corresponding authors upon reasonable request.

## Code availability

The custom scripts are available from the corresponding authors upon reasonable request.

