## Supplementary Information for "Virus-like particle delivery enables orthogonal genome editing *in vitro* and *in vivo*"

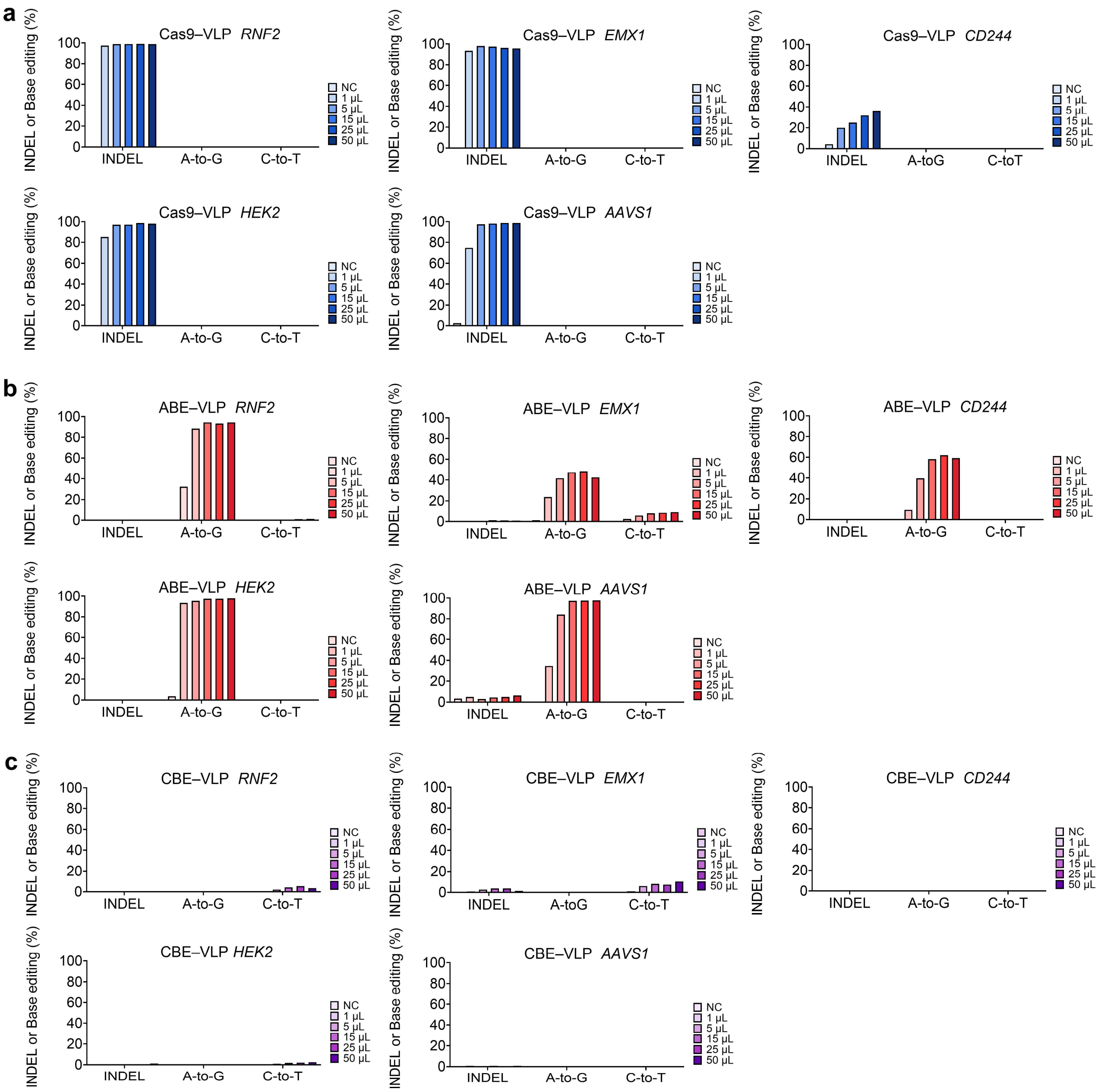


**Supplementary Fig. 1. Editing efficiencies of virus-like particles at selected genomic loci. a**, Editing efficiencies of Cas9–virus-like particle (VLP) delivery at the *RNF2*, *EMX1*, *CD244*, *HEK2*, and *AAVS1* loci in HEK293T cells. **b**, Editing efficiencies of ABE–VLP delivery at the *RNF2*, *EMX1*, *CD244*, *HEK2*, and *AAVS1* loci. **c**, Editing efficiencies of CBE-VLP delivery at the *RNF2*, *EMX1*, *CD244*, *HEK2*, and *AAVS1* loci.

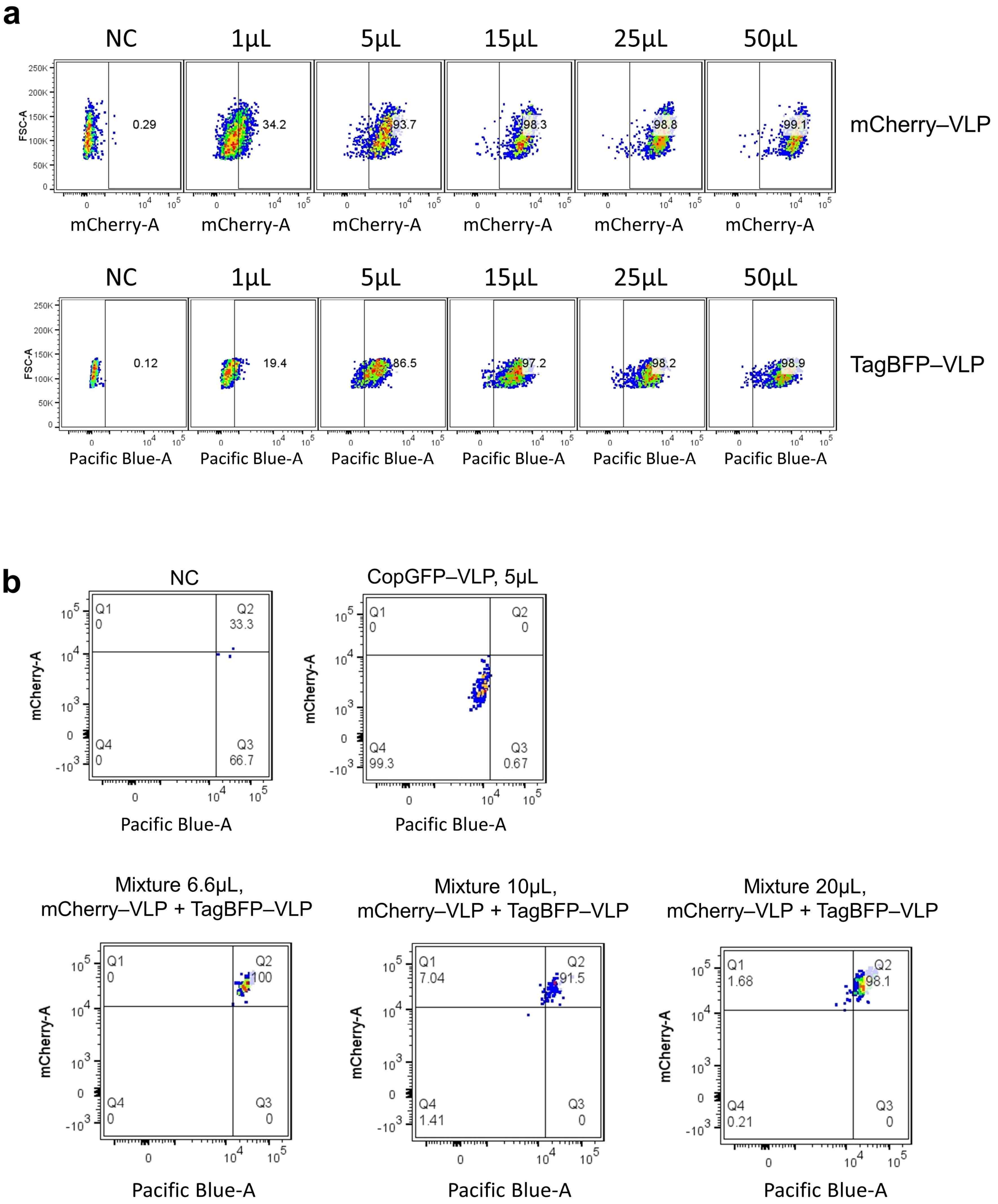


**Supplementary Fig. 2. Independent and co-delivery of mCherry– and TagBFP–VLPs. a**, Flow cytometry results of mCherry–VLP and TagBFP–VLP independent delivery in HEK293T cells. **b**, Flow cytometry results of mCherry–VLP and TagBFP–VLP co-delivery.

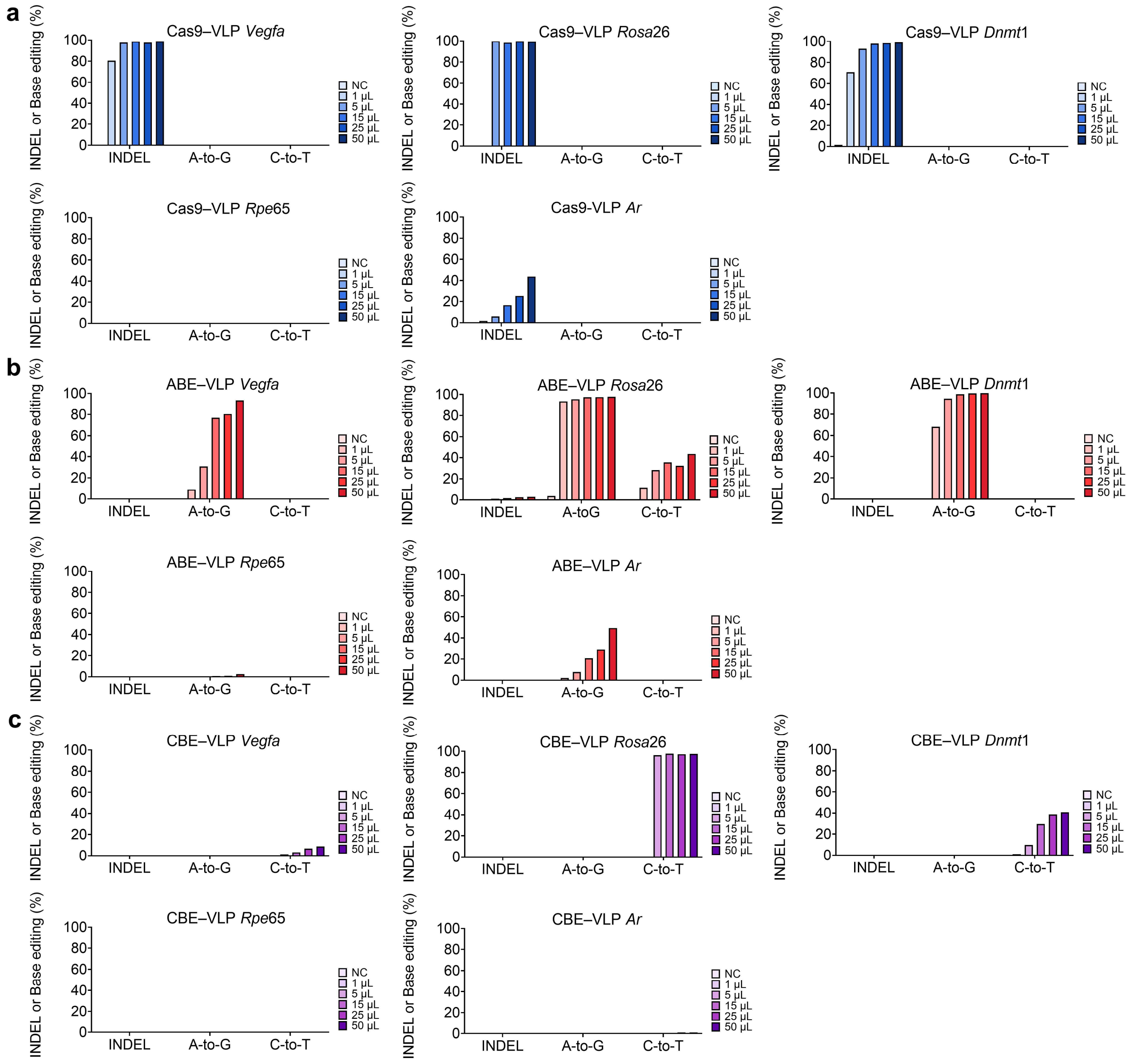


**Supplementary Fig. 3. Editing efficiencies of VLPs in mouse cells. a**, Editing efficiencies for the Cas9–VLP delivery at the *Vegfa*, *Rosa26*, *Dnmt1*, *Rpe65* and *Ar* genomic loci in NIH3T3 cells. **b**, Editing efficiencies for the ABE–VLP delivery at the *Vegfa*, *Rosa26*, *Dnmt1*, *Rpe65* and *Ar* genomic loci. **c**, Editing efficiencies for the CBE–VLP delivery at the *Vegfa*, *Rosa26*, *Dnmt1*, *Rpe65* and *Ar* genomic loci.

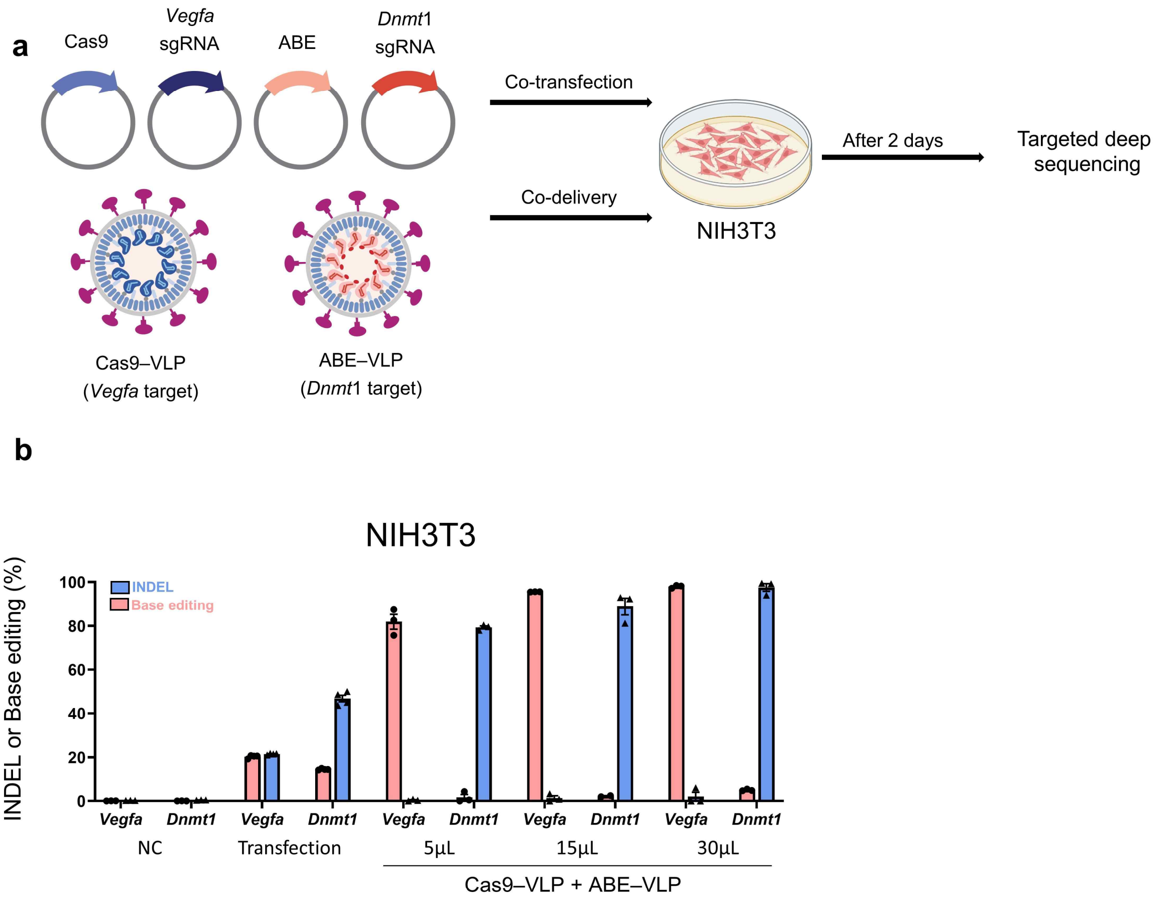


**Supplementary Fig. 4. Orthogonal editing in mouse cells using VLPs. a**, Schematic of plasmid co-transfection and VLP co-delivery at the *Vegfa* and *Dnmt1* genomic loci, followed by sequencing two days post-injection. **b**, Editing efficiencies of plasmid transfection and VLP delivery at the *Vegfa* and *Dnmt1* genomic loci in NIH3T3 cells.

**Supplementary Table 1. Plasmid sequences.**

**pGag-SpCas9**

MMLVgag – 3NES – SV40NLS – SpCas9 – SV40NLS – Myc NLS - β-globin poly(A) signal

atgggccagactgttaccactcccttaagtttgaccttaggtcactggaaagatgtcgagcggatcgctcacaaccagtcggtagatgtcaagaagagacgttgggttaccttctgctctgcagaatggccaacctttaacgtcggatggccg
cgagacggcacctttaaccgagacctcatcacccaggttaagatcaaggtcttttcacctggcccgcatggacacccagaccaggtcccctacatcgtgacctgggaagccttggcttttgacccccctccctgggtcaagccctttgtacac
cctaagcctccgcctcctcttcctccatccgccccgtctctcccccttgaacctcctcgttcgaccccgcctcgatcctccctttatccagccctcactccttctctaggcgccaaacctaaacctcaagttctttctgacagtggggggccgctcatc
gacctacttacagaagacccccgccttatagggacccaagaccaccccccttccgacagggacggaaatggtggagaagcgacccctgcgggagaggcaccggacccctccccaatggcatctcgcctacgtgggagacgggag
ccccctgtggccgactccactacctcgcaggcattccccctccgcgcaggaggaaacggacagcttcaatactggccgttctcctcttctgacctttacaactggaaaaataataacccttctttttctgaagatccaggtaaactgacagctct
gatcgagtctgtcctcatcacccatcagcccacctgggacgactgtcagcagctgttggggactctgctgaccggagaagaaaaacaacgggtgctcttagaggctagaaaggcggtgcggggcgatgatgggcgccccactcaactg
cccaatgaagtcgatgccgcttttcccctcgagcgcccagactgggattacaccacccaggcaggtaggaaccacctagtccactatcgccagttgctcctagcgggtctccaaaacgcgggcagaagccccaccaatttggccaaggt
aaaaggaataacacaagggcccaatgagtctccctcggccttcctagagagacttaaggaagcctatcgcaggtacactccttatgaccctgaggacccagggcaagaaactaatgtgtctatgtctttcatttggcagtctgccccagac
attgggagaaagttagagaggttagaagatttaaaaaacaagacgcttggagatttggttagagaggcagaaaagatctttaataaacgagaaaccccggaagaaagagaggaacgtatcaggagagaaacagaggaaaaagaa
gaacgccgtaggacagaggatgagcagaaagagaaagaaagagatcgtaggagacatagagagatgagcaagctattggccactgtcgttagtggacagaaacaggatagacagggaggagaacgaaggaggtcccaactc
gatcgcgaccagtgtgcctactgcaaagaaaaggggcactgggctaaagattgtcccaagaaaccacgaggacctcggggaccaagaccccagacctccctcctgaccctagatgactctggcggctcacttcaactgcctccacttg
aaagactgacactgggatcattacaattacctcctttagaacgattaacactcggttcactacagcttccgcctcttgagagattgacattaacgtccacgctgctaatggagaactcgtctggagattacaaagacgatgacgataagaaac
ggacagccgacggaagcgagttcgagtcaccaaagaagaagcggaaagtcGGagacaagaagtacagcatcggcctggacatcggtaccaacagcgtgggctgggccgtgatcaccgacgagtacaaggtgcccagcaaga
agttcaaggtgctgggcaacaccgaccgccacagcatcaagaagaacctgatcggcgccctgctgttcgacagcggcgagaccgccgaggccacccgcctgaagcgcaccgcccgccgccgctacacccgccgcaagaaccgc
atctgctacctgcaggagatcttcagcaacgagatggccaaggtggacgacagcttcttccaccgcctggaggagagcttcctggtggaggaggacaagaagcacgagcgccaccccatcttcggcaacatcgtggacgaggtggcc
taccacgagaagtaccccaccatctaccacctgcgcaagaagctggtggacagcaccgacaaggccgacctgcgcctgatctacctggccctggcccacatgatcaagttccgcggccacttcctgatcgagggcgacctgaacccc
gacaacagcgacgtggacaagctgttcatccagctggtgcagacctacaaccagctgttcgaggagaaccccatcaacgccagcggcgtggacgccaaggccatcctgagcgcccgcctgagcaagagccgccgcctggagaac
ctgatcgcccagctgcccggcgagaagaagaacggcctgttcggcaacctgatcgccctgagcctgggcctgacccccaacttcaagagcaacttcgacctggccgaggacgccaagctgcagctgagcaaggacacctacgacg
acgacctggacaacctgctggcccagatcggcgaccagtacgccgacctgttcctggccgccaagaacctgagcgacgccatcctgctgagcgacatcctgcgcgtgaacaccgagatcaccaaggcccccctgagcgccagcatg
atcaagcgctacgacgagcaccaccaggacctgaccctgctgaaggccctggtgcgccagcagctgcccgagaagtacaaggagatcttcttcgaccagagcaagaacggctacgccggctacatcgacggcggcgccagccag
gaggagttctacaagttcatcaagcccatcctggagaagatggacggcaccgaggagctgctggtgaagctgaaccgcgaggacctgctgcgcaagcagcgcaccttcgacaacggcagcatcccccaccagatccacctgggcg
agctgcacgccatcctgcgccgccaggaggacttctaccccttcctgaaggacaaccgcgagaagatcgagaagatcctgaccttccgcatcccctactacgtgggccccctggcccgcggcaacagccgcttcgcctggatgacccg
caagagcgaggagaccatcacccccctggaacttcgaggaggtggtggacaagggcgccagcgcccagagcttcatcgagcgcatgaccaacttcgacaagaacctgcccaacgagaaggtgctgcccaagcacagcctgctgta
cgagtacttcaccgtgtacaacgagctgaccaaggtgaagtacgtgaccgagggcatgcgcaagcccgccttcctgagcggcgagcagaagaaggccatcgtggacctgctgttcaagaccaaccgcaaggtgaccgtgaagcag
ctgaaggaggactacttcaagaagatcgagtgcttcgacagcgtggagatcagcggcgtggaggaccgcttcaacgccagcctgggcacctaccacgacctgctgaagatcatcaaggacaaggacttcctggacaacgaggagaa
cgaggacatcctggaggacatcgtgctgaccctgaccctgttcgaggaccgcgagatgatcgaggagcgcctgaagacctacgcccacctgttcgacgacaaggtgatgaagcagctgaagcgccgccgctacaccggctggggcc
gcctgagccgcaagcttatcaacggcatccgcgacaagcagagcggcaagaccatcctggacttcctgaagagcgacggcttcgccaaccgcaacttcatgcagctgatccacgacgacagcctgaccttcaaggaggacatccag
aaggcccaggtgagcggccagggcgacagcctgcacgagcacatcgccaacctggccggcagccccgccatcaagaagggcatcctgcagaccgtgaaggtggtggacgagctggtgaaggtgatgggccgccacaagcccg
agaacatcgtgatcgagatggcccgcgagaaccagaccacccagaagggccagaagaacagccgcgagcgcatgaagcgcatcgaggagggcatcaaggagctgggcagccagatcctgaaggagcaccccgtggagaac
acccagctgcagaacgagaagctgtacctgtactacctgcagaacggccgcgacatgtacgtggaccaggagctggacatcaaccgcctgagcgactacgacgtggaccacatcgtgccccagagcttcctgaaggacgacagcat
cgacaacaaggtgctgacccgcagcgacaagaaccgcggcaagagcgacaacgtgcccagcgaggaggtggtgaagaagatgaagaactactggcgccagctgctgaacgccaagctgatcacccagcgcaagttcgacaac
ctgaccaaggccgagcgcggcggcctgagcgagctggacaaggccggcttcatcaagcgccagctggtggagacccgccagatcaccaagcacgtggcccagatcctggacagccgcatgaacaccaagtacgacgagaacg
acaagctgatccgcgaggtgaaggtgatcaccctgaagagcaagctggtgagcgacttccgcaaggacttccagttctacaaggtgcgcgagatcaacaactaccaccacgcccacgacgcctacctgaacgccgtggtgggcacc
gccctgatcaagaagtaccccaagctggagagcgagttcgtgtacggcgactacaaggtgtacgacgtgcgcaagatgatcgccaagagcgagcaggagatcggcaaggccaccgccaagtacttcttctacagcaacatcatgaa
cttcttcaagaccgagatcaccctggccaacggcgagatccgcaagcgcccccctgatcgagaccaacggcgagaccggcgagatcgtgtgggacaagggccgcgacttcgccaccgtgcgcaaggtgctgagcatgccccaggtg
aacatcgtgaagaagaccgaggtgcagaccggcggcttcagcaaggagagcatcctgcccaagcgcaacagcgacaagctgatcgcccgcaagaaggactgggaccccaagaagtacggcggcttcgacagccccaccgtgg
cctacagcgtgctggtggtggccaaggtggagaagggcaagagcaagaagctgaagagcgtgaaggagctgctgggcatcaccatcatggagcgcagcagcttcgagaagaaccccatcgacttcctggaggccaagggctaca
aggaggtgaagaaggacctgatcatcaagctgcccaagtacagcctgttcgagctggagaacggccgcaagcgcatgctggccagcgccggcgagctgcagaagggcaacgagctggccctgcccagcaagtacgtgaacttcct
gtacctggccagccactacgagaagctgaagggcagccccgaggacaacgagcagaagcagctgttcgtggagcagcacaagcactacctggacgagatcatcgagcagatcagcgagttcagcaagcgcgtgatcctggccga
cgccaacctggacaaggtgctgagcgcctacaacaagcaccgcgacaagcccatccgcgagcaggccgagaacatcatccacctgttcaccctgaccaacctgggcgcccccgccgccttcaagtacttcgacaccaccatcgacc
gcaagcgctacaccagcaccaaggaggtgctggacgccaccctgatccaccagagcatcaccggtctgtacgagacccgcatcgacctgagccagctgggcggcgactctggcggctcaaaaagaaccgccgacggcagcgaat
tcgagtctcccaagaagaagaggaaagtcggctctggccctgccgctaagagagtgaagctggactgagatctttttccctctgccaaaaattatggggacatcatgaagccccttgagcatctgacttctggctaataaaggaaatttatttt
cattgcaatagtgtgttggaattttttgtgtctctca

**pGag-ABE8e**

MMLVgag – 3NES – SV40NLS – TadA 8e nCas9 (D10A) – SV40NLS – Myc NLS - β-globin poly(A) signal

atgggccagactgttaccactcccttaagtttgaccttaggtcactggaaagatgtcgagcggatcgctcacaaccagtcggtagatgtcaagaagagacgttgggttaccttctgctctgcagaatggccaacctttaacgtcggatggccg
cgagacggcacctttaaccgagacctcatcacccaggttaagatcaaggtcttttcacctggcccgcatggacacccagaccaggtcccctacatcgtgacctgggaagccttggcttttgacccccctccctgggtcaagccctttgtacac
cctaagcctccgcctcctcttcctccatccgccccgtctctcccccttgaacctcctcgttcgaccccgcctcgatcctccctttatccagccctcactccttctctaggcgccaaacctaaacctcaagttctttctgacagtggggggccgctcatc
gacctacttacagaagacccccgccttatagggacccaagaccaccccccttccgacagggacggaaatggtggagaagcgacccctgcgggagaggcaccggacccctccccaatggcatctcgcctacgtgggagacgggag
ccccctgtggccgactccactacctcgcaggcattccccctccgcgcaggaggaaacggacagcttcaatactggccgttctcctcttctgacctttacaactggaaaaataataacccttctttttctgaagatccaggtaaactgacagctct
gatcgagtctgtcctcatcacccatcagcccacctgggacgactgtcagcagctgttggggactctgctgaccggagaagaaaaacaacgggtgctcttagaggctagaaaggcggtgcggggcgatgatgggcgccccactcaactg
cccaatgaagtcgatgccgcttttcccctcgagcgcccagactgggattacaccacccaggcaggtaggaaccacctagtccactatcgccagttgctcctagcgggtctccaaaacgcgggcagaagccccaccaatttggccaaggt
aaaaggaataacacaagggcccaatgagtctccctcggccttcctagagagacttaaggaagcctatcgcaggtacactccttatgaccctgaggacccagggcaagaaactaatgtgtctatgtctttcatttggcagtctgccccagac
attgggagaaagttagagaggttagaagatttaaaaaacaagacgcttggagatttggttagagaggcagaaaagatctttaataaacgagaaaccccggaagaaagagaggaacgtatcaggagagaaacagaggaaaaagaa
gaacgccgtaggacagaggatgagcagaaagagaaagaaagagatcgtaggagacatagagagatgagcaagctattggccactgtcgttagtggacagaaacaggatagacagggaggagaacgaaggaggtcccaactc
gatcgcgaccagtgtgcctactgcaaagaaaaggggcactgggctaaagattgtcccaagaaaccacgaggacctcggggaccaagaccccagacctccctcctgaccctagatgactctggcggctcacttcaactgcctccacttg
aaagactgacactgggatcattacaattacctcctttagaacgattaacactcggttcactacagcttccgcctcttgagagattgacattaacgtccacgctgctaatggagaactcgtctggagattacaaagacgatgacgataagaaac
ggacagccgacggaagcgagttcgagtcaccaaagaagaagcggaaagtcGGatctgaggtggagttttcccacgagtactggatgagacatgccctgaccctggccaagagggcacgggatgagagggaggtgcctgtggga
gccgtgctggtgctgaacaatagagtgatcggcgagggctggaacagagccatcggcctgcacgacccaacagcccatgccgaaattatggccctgagacagggcggcctggtcatgcagaactacagactgattgacgccaccctg
tacgtgacattcgagccttgcgtgatgtgcgccggcgccatgatccactctaggatcggccgcgtggtgtttggcgtgaggaactcaaaaagaggcgccgcaggctccctgatgaacgtgctgaactaccccggcatgaatcaccgcgtc
gaaattaccgagggaatcctggcagatgaatgtgccgccctgctgtgcgatttctatcggatgcctagacaggtgttcaatgctcagaagaaggcccagagctccatcaactccggaggatctagcggaggctcctctggctctgagacac
ctggcacaagcgagagcgcaacacctgaaagcagcgggggcagcagcggggggtcagacaagaagtacagcatcggcctggccatcggcaccaactctgtgggctgggccgtgatcaccgacgagtacaaggtgcccagcaa
gaaattcaaggtgctgggcaacaccgaccggcacagcatcaagaagaacctgatcggagccctgctgttcgacagcggcgaaacagccgaggccacccggctgaagagaaccgccagaagaagatacaccagacggaagaa
ccggatctgctatctgcaagagatcttcagcaacgagatggccaaggtggacgacagcttcttccacagactggaagagtccttcctggtggaagaggataagaagcacgagcggcaccccatcttcggcaacatcgtggacgaggtg
gcctaccacgagaagtaccccaccatctaccacctgagaaagaaactggtggacagcaccgacaaggccgacctgcggctgatctatctggccctggcccacatgatcaagttccggggccacttcctgatcgagggcgacctgaac
cccgacaacagcgacgtggacaagctgttcatccagctggtgcagacctacaaccagctgttcgaggaaaaccccatcaacgccagcggcgtggacgccaaggccatcctgtctgccagactgagcaagagcagacggctggaaa
atctgatcgcccagctgcccggcgagaagaagaatggcctgttcggaaacctgattgccctgagcctgggcctgacccccaacttcaagagcaacttcgacctggccgaggatgccaaactgcagctgagcaaggacacctacgacg
acgacctggacaacctgctggcccagatcggcgaccagtacgccgacctgtttctggccgccaagaacctgtccgacgccatcctgctgagcgacatcctgagagtgaacaccgagatcaccaaggcccccctgagcgcctctatgat
caagagatacgacgagcaccaccaggacctgaccctgctgaaagctctcgtgcggcagcagctgcctgagaagtacaaagagattttcttcgaccagagcaagaacggctacgccggctacattgacggcggagccagccaggaa
gagttctacaagttcatcaagcccatcctggaaaagatggacggcaccgaggaactgctcgtgaagctgaacagagaggacctgctgcggaagcagcggaccttcgacaacggcagcatcccccaccagatccacctgggagagct
gcacgccattctgcggcggcaggaagatttttacccattcctgaaggacaaccgggaaaagatcgagaagatcctgaccttccgcatcccctactacgtgggccctctggccagggaaacagcagattcgcctggatgaccagaaag
agcgaggaaaccatcaccccctggaacttcgaggaagtggtggacaagggcgcttccgcccagagcttcatcgagcggatgaccaacttcgataagaacctgcccaacgagaaggtgctgcccaagcacagcctgctgtacgagta
cttcaccgtgtataacgagctgaccaaagtgaaatacgtgaccgagggaatgagaaagcccgccttcctgagcggcgagcagaaaaaggccatcgtggacctgctgttcaagaccaaccggaaagtgaccgtgaagcagctgaaa
gaggactacttcaagaaaatcgagtgcttcgactccgtggaaatctccggcgtggaagatcggttcaacgcctccctgggcacataccacgatctgctgaaaattatcaaggacaaggacttcctggacaatgaggaaaacgaggacat
tctggaagatatcgtgctgaccctgacactgtttgaggacagagagatgatcgaggaacggctgaaaacctatgcccacctgttcgacgacaaagtgatgaagcagctgaagcggcggagatacaccggctggggcaggctgagccg
gaagctgatcaacggcatccgggacaagcagtccggcaagacaatcctggatttcctgaagtccgacggcttcgccaacagaaacttcatgcagctgatccacgacgacagcctgacctttaaagaggacatccagaaagcccaggt
gtccggccagggcgatagcctgcacgagcacattgccaatctggccggcagccccgccattaagaagggcatcctgcagacagtgaaggtggtggacgagctcgtgaaagtgatgggccggcacaagcccgagaacatcgtgatc
gaaatggccagagagaaccagaccacccagaagggacagaagaacagccgcgagagaatgaagcggatcgaagagggcatcaaagagctgggcagccagatcctgaaagaacaccccgtggaaaacacccagctgcaga
acgagaagctgtacctgtactacctgcagaatgggcgggatatgtacgtggaccaggaactggacatcaaccggctgtccgactacgatgtggaccatatcgtgcctcagagctttctgaaggacgactccatcgacaacaaggtgctga
ccagaagcgacaagaaccggggcaagagcgacaacgtgccctccgaagaggtcgtgaagaagatgaagaactactggcggcagctgctgaacgccaagctgattacccagagaaagttcgacaatctgaccaaggccgagag
aggcggcctgagcgaactggataaggccggcttcatcaagagacagctggtggaaacccggcagatcacaaagcacgtggcacagatcctggactcccggatgaacactaagtacgacgagaatgacaagctgatccgggaagt
gaaagtgatcaccctgaagtccaagctggtgtccgatttccggaaggatttccagttttacaaagtgcgcgagatcaacaactaccaccacgcccacgacgcctacctgaacgccgtcgtgggaaccgccctgatcaaaaagtaccct
aagctggaaagcgagttcgtgtacggcgactacaaggtgtacgacgtgcggaagatgatcgccaagagcgagcaggaaatcggcaaggctaccgccaagtacttcttctacagcaacatcatgaactttttcaagaccgagattaccctg
gccaacggcgagatccggaagcggcctctgatcgagacaaacggcgaaaccggggagatcgtgtgggataagggccgggattttgccaccgtgcggaaagtgctgagcatgccccaagtgaatatcgtgaaaaagaccgaggtgc
agacaggcggcttcagcaaagagtctatcctgcccaagaggaacagcgataagctgatcgccagaaagaaggactgggaccctaagaagtacggcggcttcgacagccccaccgtggcctattctgtgctggtggtggccaaagtgg
aaaagggcaagtccaagaaactgaagagtgtgaaagagctgctggggatcaccatcatggaaagaagcagcttcgagaagaatcccatcgactttctggaagccaagggctacaaagaagtgaaaaaggacctgatcatcaagct
gcctaagtactccctgttcgagctggaaaacggccggaagagaatgctggcctctgccggcgaactgcagaagggaaacgaactggccctgccctccaaatatgtgaacttcctgtacctggccagccactatgagaagctgaagggc
tcccccgaggataatgagcagaaacagctgtttgtggaacagcacaagcactacctggacgagatcatcgagcagatcagcgagttctccaagagagtgatcctggccgacgctaatctggacaaagtgctgtccgcctacaacaagc
accgggataagcccatcagagagcaggccgagaatatcatccacctgtttaccctgaccaatctgggagcccctgccgccttcaagtactttgacaccaccatcgaccggaagaggtacaccagcaccaaagaggtgctggacgcca
ccctgatccaccagagcatcaccggcctgtacgagacacggatcgacctgtctcagctgggaggtgactctggcggctcaaaaagaaccgccgacggcagcgaattcgagtctcccaagaagaagaggaaagtcggctctggccct
gccgctaagagagtgaagctggactgagatctttttccctctgccaaaaattatggggacatcatgaagccccttgagcatctgacttctggctaataaaggaaatttattttcattgcaatagtgtgttggaattttttgtgtctctca

**pGag-AncBE4max**

MMLVgag – 3NES – SV40NLS – AncAPOBEC nCas9 (D10A) UGI – SV40NLS – Myc NLS - β-globin poly(A) signal

atgggccagactgttaccactcccttaagtttgaccttaggtcactggaaagatgtcgagcggatcgctcacaaccagtcggtagatgtcaagaagagacgttgggttaccttctgctctgcagaatggccaacctttaacgtcggatggccg
cgagacggcacctttaaccgagacctcatcacccaggttaagatcaaggtcttttcacctggcccgcatggacacccagaccaggtcccctacatcgtgacctgggaagccttggcttttgacccccctccctgggtcaagccctttgtacac
cctaagcctccgcctcctcttcctccatccgccccgtctctcccccttgaacctcctcgttcgaccccgcctcgatcctccctttatccagccctcactccttctctaggcgccaaacctaaacctcaagttctttctgacagtgggggccgctcatc
gacctacttacagaagacccccgccttatagggacccaagaccacccccttccgacagggacggaaatggtggagaagcgacccctgcgggagaggcaccggacccctccccaatggcatctcgcctacgtgggagacgggag
ccccctgtggccgactccactacctcgcaggcattccccctccgcgcaggaggaaacggacagcttcaatactggccgttctcctcttctgacctttacaactggaaaaataataacccttctttttctgaagatccaggtaaactgacagctct
gatcgagtctgtcctcatcacccatcagcccacctgggacgactgtcagcagctgttggggactctgctgaccggagaagaaaaacaacgggtgctcttagaggctagaaaggcggtgcggggcgatgatgggcgccccactcaactg
cccaatgaagtcgatgccgcttttccccctcgagcgcccagactgggattacaccacccaggcaggtaggaaccacctagtccactatcgccagttgctcctagcgggtctccaaaacgcgggcagaagccccaccaatttggccaaggt
aaaaggaataacacaagggcccaatgagtctccctcggccttcctagagagacttaaggaagcctatcgcaggtacactccttatgaccctgaggacccagggcaagaaactaatgtgtctatgtctttcatttggcagtctgccccagac
attgggagaaagttagagaggttagaagatttaaaaaacaagacgcttggagatttggttagagaggcagaaaagatctttaataaacgagaaaccccggaagaaagagaggaacgtatcaggagagaaacagaggaaaaagaa
gaacgccgtaggacagaggatgagcagaaagagaaagaaagagatcgtaggagacatagagagatgagcaagctattggccactgtcgttagtggacagaaacaggatagacagggaggagaacgaaggaggtcccaactc
gatcgcgaccagtgtgcctactgcaaagaaaaggggcactgggctaaagattgtcccaagaaaccacgaggacctcggggaccaagaccccagacctccctcctgaccctagatgactctggcggctcacttcaactgcctccacttg
aaagactgacactgggatcattacaattacctccttagaacgattaacactcggttcactacagcttccgcctcttgagagattgacattaacgtccacgctgctaatggagaactcgtctggagattacaaagacgatgacgataagaaac
ggacagccgacggaagcgagttcgagtcaccaaagaagaagcggaaagtcGGaagcagtgaaaccggaccagtggcagtggacccaaccctgaggagacggattgagccccatgaatttgaagtgttctttgacccaagggag
ctgaggaaggagacatgcctgctgtacgagatcaagtggggcacaagccacaagatctggcgccacagctccaagaacaccacaaagcacgtggaagtgaatttcatcgagaagtttacctccgagcggcacttctgcccctctacca
gctgttccatcacatggtttctgtcttggagccctgcggcgagtgttccaaggccatcaccgagttcctgtctcagcaccctaacgtgaccctggtcatctacgtggcccggctgtatcaccacatggaccagcagaacaggcagggcctgc
gcgatctggtgaattctggcgtgaccatccagatcatgacagccccagagtacgactattgctggcggaacttcgtgaattatccacctggcaaggaggcacactggccaagatacccacccctgtggatgaagctgtatgcactggagct
gcacgcaggaatcctgggcctgcctccatgtctgaatatcctgcggagaaagcagccccagctgacatttttcaccattgctctgcagtcttgtcactatcagcggctgcctcctcatattctgtgggctacaggcctgaagtctggaggatcta
gcggaggatcctctggcagcgagacaccaggaacaagcgagtcagcaacaccagagagcagtggcggcagcagcggcggcagcgacaagaagtacagcatcggcctggccatcggcaccaactctgtgggctgggccgtgat
caccgacgagtacaaggtgcccagcaagaaattcaaggtgctgggcaacaccgaccggcacagcatcaagaagaacctgatcggagccctgctgttcgacagcggcgaaacagccgaggccacccggctgaagagaaccgcc
agaagaagatacaccagacggaagaaccggatctgctatctgcaagagatcttcagcaacgagatggccaaggtggacgacagcttcttccacagactggaagagtccttcctggtggaagaggataagaagcacgagcggcacc
ccatcttcggcaacatcgtggacgaggtggcctaccacgagaagtaccccaccatctaccacctgagaaagaaactggtggacagcaccgacaaggccgacctgcggctgatctatctggccctggcccacatgatcaagttccgggg
ccacttcctgatcgagggcgacctgaaccccgacaacagcgacgtggacaagctgttcatccagctggtgcagacctacaaccagctgttcgaggaaaaccccatcaacgccagcggcgtggacgccaaggccatcctgtctgccag
actgagcaagagcagacggctggaaaatctgatcgcccagctgcccggcgagaagaagaatggcctgttcggaaacctgattgccctgagcctgggcctgacccccaacttcaagagcaacttcgacctggccgaggatgccaaact
gcagctgagcaaggacacctacgacgacgacctggacaacctgctggcccagatcggcgaccagtacgccgacctgtttctggccgccaagaacctgtccgacgccatcctgctgagcgacatcctgagagtgaacaccgagatca
ccaaggcccccctgagcgcctctatgatcaagagatacgacgagcaccaccaggacctgaccctgctgaaagctctcgtgcggcagcagctgcctgagaagtacaaagagattttcttcgaccagagcaagaacggctacgccggct
acattgacggcggagccagccaggaagagttctacaagttcatcaagcccatcctggaaaagatggacggcaccgaggaactgctcgtgaagctgaacagagaggacctgctgcggaagcagcggaccttcgacaacggcagcat
cccccaccagatccacctgggagagctgcacgccattctgcggcggcaggaagatttttacccattcctgaaggacaaccgggaaaagatcgagaagatcctgaccttccgcatcccctactacgtgggccctctggccaggggaaac
agcagattcgcctggatgaccagaaagagcgaggaaaccatcaccccctggaacttcgaggaagtggtggacaagggcgcttccgcccagagcttcatcgagcggatgaccaacttcgataagaacctgcccaacgagaaggtgct
gcccaagcacagcctgctgtacgagtacttcaccgtgtataacgagctgaccaaagtgaaatacgtgaccgagggaatgagaaagcccgccttcctgagcggcgagcagaaaaaggccatcgtggacctgctgttcaagaccaacc
ggaaagtgaccgtgaagcagctgaaagaggactacttcaagaaaatcgagtgcttcgactccgtggaaatctccggcgtggaagatcggttcaacgcctccctgggcacataccacgatctgctgaaaattatcaaggacaaggacttc
ctggacaatgaggaaaacgaggacattctggaagatatcgtgctgaccctgacactgtttgaggacagagagatgatcgaggaacggctgaaaacctatgcccacctgttcgacgacaaagtgatgaagcagctgaagcggcggag
atacaccggctggggcaggctgagccggaagctgatcaacggcatccgggacaagcagtccggcaagacaatcctggatttcctgaagtccgacggcttcgccaacagaaacttcatgcagctgatccacgacgacagcctgacctttt
aaagaggacatccagaaagcccaggtgtccggccagggcgatagcctgcacgagcacattgccaatctggccggcagccccgccattaagaagggcatcctgcagacagtgaaggtggtggacgagctcgtgaaagtgatgggcc
ggcacaagcccgagaacatcgtgatcgaaatggccagagagaaccagaccacccagaagggacagaagaacagccgcgagagaatgaagcggatcgaagagggcatcaaagagctgggcagccagatcctgaaagaaca
ccccgtggaaaacacccagctgcagaacgagaagctgtacctgtactacctgcagaatgggcgggatatgtacgtggaccaggaactggacatcaaccggctgtccgactacgatgtggaccatatcgtgcctcagagctttctgaagg
acgactccatcgacaacaaggtgctgaccagaagcgacaagaaccggggcaagagcgacaacgtgccctccgaagaggtcgtgaagaagatgaagaactactggcggcagctgctgaacgccaagctgattacccagagaaa
gttcgacaatctgaccaaggccgagagaggcggcctgagcgaactggataaggccggcttcatcaagagacagctggtggaaacccggcagatcacaaagcacgtggcacagatcctggactcccggatgaacactaagtacgac
gagaatgacaagctgatccgggaagtgaaagtgatcaccctgaagtccaagctggtgtccgatttccggaaggatttccagttttacaaagtgcgcgagatcaacaactaccaccacgcccacgacgcctacctaaacgccgtcgtggg
aaccgccctgatcaaaaagtaccctaagctggaaagcgagttcgtgtacggcgactacaaggtgtacgacgtgcggaagatgatcgccaagagcgagcaggaaatcggcaaggctaccgccaagtacttcttctacagcaacatcat
gaactttttcaagaccgagattaccctggccaacggcgagatccggaagcggcctctgatcgagacaaacggcgaaaccggggagatcgtgtgggataagggccgggattttgccaccgtgcggaaagtgctgagcatgccccaagt
gaatatcgtgaaaaagaccgaggtgcagacaggcggcttcagcaaagagtctatcctgcccaagaggaacagcgataagctgatcgccagaaagaaggactgggaccctaagaagtacggcggcttcgacagccccaccgtggc
ctattctgtgctggtggtggccaaagtggaaaagggcaagtccaagaaactgaagagtgtgaaagagctgctggggatcaccatcatggaaagaagcagcttcgagaagaatcccatcgactttctggaagccaagggctacaaaga
agtgaaaaaggacctgatcatcaagctgcctaagtactccctgttcgagctggaaaacggccggaagagaatgctggcctctgccggcgaactgcagaagggaaacgaactggccctgccctccaaatatgtgaacttcctgtacctgg
ccagccactatgagaagctgaagggctcccccgaggataatgagcagaaacagctgtttgtggaacagcacaagcactacctggacgagatcatcgagcagatcagcgagttctccaagagagtgatcctggccgacgctaatctgg
acaaagtgctgtccgcctacaacaagcaccgggataagcccatcagagagcaggccgagaatatcatccacctgtttaccctgaccaatctgggagcccctgccgccttcaagtactttgacaccaccatcgaccggaagaggtacac
cagcaccaaagaggtgctggacgccaccctgatccaccagagcatcaccggcctgtacgagacacggatcgacctgtctcagctgggaggtgacagcggcgggagcggcgggagcgggggggagcactaatctgagcgacatcat
tgagaaggagactgggaaacagctggtcattcaggagtccatcctgatgctgcctgaggaggtggaggaagtgatcggcaacaagccagagtctgacatcctggtgcacaccgcctacgacgagtccacagatgagaatgtgatgctg
ctgacctctgacgcccccgagtataagccttgggccctggtcatccaggattctaacggcgagaataagatcaagatgctgagcggaggatccggaggatctggaggcagcaccaacctgtctgacatcatcgagaaggagacaggc
aagcagctggtcatccaggagagcatcctgatgctgcccgaagaagtcgaagaagtgatcggaaacaagcctgagagcgatatcctggtccataccgcctacgacgagagtaccgacgaaaatgtgatgctgctgacatccgacgcc
ccagagtataagccctgggctctggtcatccaggattccaacggagagaacaaaatcaaaatgctgtctggcggctcaaaaagaaccgccgacggcagcgaattcgagtctcccaagaagaagaggaaagtcggctctggccctgc
cgctaagagagtgaagctggactgagatctttttccctctgccaaaaattatggggacatcatgaagccccttgagcatctgacttctggctaataaaggaaatttattttcattgcaatagtgtgttggaattttttgtgtctctca

**pGag-VP64 dCas9**

MMLVgag – 3NES – SV40NLS – VP64 dCas9 (D10A, H840A) – SV40NLS – Myc NLS - β-globin poly(A) signal

atgggccagactgttaccactcccttaagtttgaccttaggtcactggaaagatgtcgagcggatcgctcacaaccagtcggtagatgtcaagaagagacgttgggttaccttctgctctgcagaatggccaacctttaacgtcggatggccg
cgagacggcacctttaaccgagacctcatcacccaggttaagatcaaggtcttttcacctggcccgcatggacacccagaccaggtcccctacatcgtgacctgggaagccttggcttttgacccccctccctgggtcaagccctttgtacac
cctaagcctccgcctcctcttcctccatccgccccgtctctcccccttgaacctcctcgttcgaccccgcctcgatcctccctttatccagccctcactccttctctaggcgccaaacctaaacctcaagttctttctgacagtgggggccgctcatc
gacctacttacagaagacccccgccttatagggacccaagaccacccccttccgacagggacggaaatggtggagaagcgacccctgcgggagaggcaccggacccctccccaatggcatctcgcctacgtgggagacgggag
ccccctgtggccgactccactacctcgcaggcattccccctccgcgcaggaggaaacggacagcttcaatactggccgttctcctcttctgacctttacaactggaaaaataataacccttctttttctgaagatccaggtaaactgacagctct
gatcgagtctgtcctcatcacccatcagcccacctgggacgactgtcagcagctgttggggactctgctgaccggagaagaaaaacaacgggtgctcttagaggctagaaaggcggtgcggggcgatgatgggcgccccactcaactg
cccaatgaagtcgatgccgcttttccccctcgagcgcccagactgggattacaccacccaggcaggtaggaaccacctagtccactatcgccagttgctcctagcgggtctccaaaacgcgggcagaagccccaccaatttggccaaggt
aaaaggaataacacaagggcccaatgagtctccctcggccttcctagagagacttaaggaagcctatcgcaggtacactccttatgaccctgaggacccagggcaagaaactaatgtgtctatgtctttcatttggcagtctgccccagac
attgggagaaagttagagaggttagaagatttaaaaaacaagacgcttggagatttggttagagaggcagaaaagatctttaataaacgagaaaccccggaagaaagagaggaacgtatcaggagagaaacagaggaaaaagaa
gaacgccgtaggacagaggatgagcagaaagagaaagaaagagatcgtaggagacatagagagatgagcaagctattggccactgtcgttagtggacagaaacaggatagacagggaggagaacgaaggaggtcccaactc
gatcgcgaccagtgtgcctactgcaaagaaaaggggcactgggctaaagattgtcccaagaaaccacgaggacctcggggaccaagaccccagacctccctcctgaccctagatgactctggcggctcacttcaactgcctccacttg
aaagactgacactgggatcattacaattacctccttagaacgattaacactcggttcactacagcttccgcctcttgagagattgacattaacgtccacgctgctaatggagaactcgtctggagattacaaagacgatgacgataagaaac
ggacagccgacggaagcgagttcgagtcaccaaagaagaagcggaaagtcGGagacaagaagtacagcatcggcctggccatcggcaccaactctgtgggctgggccgtgatcaccgacgagtacaaggtgcccagcaagaa
attcaaggtgctgggcaacaccgaccggcacagcatcaagaagaacctgatcggagccctgctgttcgacagcggcgaaacagccgaggccacccggctgaagagaaccgccagaagaagatacaccagacggaagaaccg
gatctgctatctgcaagagatcttcagcaacgagatggccaaggtggacgacagcttcttccacagactggaagagtccttcctggtggaagaggataagaagcacgagcggcaccccatcttcggcaacatcgtggacgaggtggcct
accacgagaagtaccccaccatctaccacctgagaaagaaactggtggacagcaccgacaaggccgacctgcggctgatctatctggccctggcccacatgatcaagttccggggccacttcctgatcgagggcgacctgaaccccg
acaacagcgacgtggacaagctgttcatccagctggtgcagacctacaaccagctgttcgaggaaaaccccatcaacgccagcggcgtggacgccaaggccatcctgtctgccagactgagcaagagcagacggctggaaaatctg
atcgcccagctgcccggcgagaagaagaatggcctgttcggcaacctgattgccctgagcctgggcctgacccccaacttcaagagcaacttcgacctggccgaggatgccaaactgcagctgagcaaggacacctacgacgacga
cctggacaacctgctggcccagatcggcgaccagtacgccgacctgtttctggccgccaagaacctgtccgacgccatcctgctgagcgacatcctgagagtgaacaccgagatcaccaaggcccccctgagcgcctctatgatcaag
agatacgacgagcaccaccaggacctgaccctgctgaaagctctcgtgcggcagcagctgcctgagaagtacaaagagattttcttcgaccagagcaagaacggctacgccggctacattgacggcggagccagccaggaagagtt
ctacaagttcatcaagcccatcctggaaaagatggacggcaccgaggaactgctcgtgaagctgaacagagaggacctgctgcggaagcagcggaccttcgacaacggcagcatcccccaccagatccacctgggagagctgcac
gccattctgcggcggcaggaagatttttacccattcctgaaggacaaccgggaaaagatcgagaagatcctgaccttccgcatcccctactacgtgggccctctggccaggggaaacagcagattcgcctggatgaccagaaagagcg
aggaaaccatcaccccctggaacttcgaggaagtggtggacaagggcgcttccgcccagagcttcatcgagcggatgaccaacttcgataagaacctgcccaacgagaaggtgctgcccaagcacagcctgctgtacgagtacttca
ccgtgtataacgagctgaccaaagtgaaatacgtgaccgagggaatgagaaagcccgccttcctgagcggcgagcagaaaaaggccatcgtggacctgctgttcaagaccaaccggaaagtgaccgtgaagcagctgaaagagg
actacttcaagaaaatcgagtgcttcgactccgtggaaatctccggcgtggaagatcggttcaacgcctccctgggcacataccacgatctgctgaaaattatcaaggacaaggacttcctggacaatgaggaaaacgaggacattctgg
aagatatcgtgctgaccctgacactgtttgaggacagagagatgatcgaggaacggctgaaaacctatgcccacctgttcgacgacaaagtgatgaagcagctgaagcggcggagatacaccggctggggcaggctgagccggaag
ctgatcaacggcatccgggacaagcagtccggcaagacaatcctggatttcctgaagtccgacggcttcgccaacagaaacttcatgcagctgatccacgacgacagcctgacctttaaagaggacatccagaaagcccaggtgtccg
gccagggcgatagcctgcacgagcacattgccaatctggccggcagccccgccattaagaagggcatcctgcagacagtgaaggtggtggacgagctcgtgaaagtgatgggccggcacaagcccgagaacatcgtgatcgaaat
ggccagagagaaccagaccacccagaagggacagaagaacagccgcgagagaatgaagcggatcgaagagggcatcaaagagctgggcagccagatcctgaaagaacaccccgtggaaaacacccagctgcagaacga
gaagctgtacctgtactacctgcagaatgggcgggatatgtacgtggaccaggaactggacatcaaccggctgtccgactacgatgtggaccacatcgtgcctcagagctttctgaaggacgactccatcgacaacaaggtgctgacca
gaagcgacaaggcccggggcaagagcgacaacgtgccctccgaagaggtcgtgaagaagatgaagaactactggcggcagctgctgaacgccaagctgattacccagagaaagttcgacaatctgaccaaggccgagagagg
cggcctgagcgaactggataaggccggcttcatcaagagacagctggtggaaacccggcagatcacaaagcacgtggcacagatcctggactcccggatgaacactaagtacgacgagaatgacaagctgatccgggaagtgaa
agtgatcaccctgaagtccaagctggtgtccgatttccggaaggatttccagttttacaaagtgcgcgagatcaacaactaccaccacgcccacgacgcctacctgaacgccgtcgtgggaaccgccctgatcaaaaagtaccctaagct
ggaaagcgagttcgtgtacggcgactacaaggtgtacgacgtgcggaagatgatcgccaagagcgagcaggaaatcggcaaggctaccgccaagtacttcttctacagcaacatcatgaactttttcaagaccgagattaccctggcc
aacggcgagatccggaagcggcctctgatcgagacaaacggcgaaaccggggagatcgtgtgggataagggccgggattttgccaccgtgcggaaagtgctgagcatgccccaagtgaatatcgtgaaaaagaccgaggtgcag
acaggcggcttcagcaaagagtctatcctgcccaagaggaacagcgataagctgatcgccagaaagaaggactgggaccctaagaagtacggcggcttcgacagccccaccgtggcctattctgtgctggtggtggccaaagtggaa
aagggcaagtccaagaaactgaagagtgtgaaagagctgctggggatcaccatcatggaaagaagcagcttcgagaagaatcccatcgactttctggaagccaagggctacaaagaagtgaaaaaggacctgatcatcaagctgc
ctaagtactccctgttcgagctggaaaacggccggaagagaatgctggcctctgccggcgaactgcagaagggaaacgaactggccctgccctccaaatatgtgaacttcctgtacctggccagccactatgagaagctgaagggctcc
cccgaggataatgagcagaaacagctgtttgtggaacagcacaagcactacctggacgagatcatcgagcagatcagcgagttctccaagagagtgatcctggccgacgctaatctggacaaagtgctgtccgcctacaacaagcac
cgggataagcccatcagagagcaggccgagaatatcatccacctgtttaccctgaccaatctgggagcccctgccgccttcaagtactttgacaccaccatcgaccggaagaggtacaccagcaccaaagaggtgctggacgccacc
ctgatccaccagagcatcaccggcctgtacgagacacggatcgacctgtctcagctgggaggcgacagcgctggaggaggtggaagcggaggaggaggaagcggaggaggaggtagcggacctaagaaaaagaggaaggtg
gcggccgctggatccggacgggctgacgcattggacgattttgatctggatatgctgggaagtgacgccctcgatgattttgaccttgacatgcttggttcggatgcccttgatgactttgacctcgacatgctcggcagtgacgcccttgatgatt
tcgacctggacatgctgtctggcggctcaaaaagaaccgccgacggcagcgaattcgagtctcccaagaagaagaggaaagtcggctctggccctgccgctaagagagtgaagctggactgagatctttttccctctgccaaaaattatg
gggacatcatgaagccccttgagcatctgacttctggctaataaaggaaatttattttcattgcaatagtgtgttggaattttttgtgtctctca

**pGag-KRAB dCas9**

MMLVgag – 3NES – SV40NLS – KRAB dCas9 (D10A, H840A) – SV40NLS – Myc NLS - β-globin poly(A) signal

atgggccagactgttaccactcccttaagtttgaccttaggtcactggaaagatgtcgagcggatcgctcacaaccagtcggtagatgtcaagaagagacgttgggttaccttctgctctgcagaatggccaacctttaacgtcggatggccg
cgagacggcacctttaaccgagacctcatcacccaggttaagatcaaggtcttttcacctggcccgcatggacacccagaccaggtccccctacatcgtgacctgggaagccttggcttttgaccccccctccctgggtcaagccctttgtacac
cctaagcctccgcctcctcttcctccatccgccccgtctctcccccttgaacctcctcgttcgaccccgcctcgatcctccctttatccagccctcactccttctctaggcgccaaacctaaacctcaagttctttctgacagtgggggccgctcatc
gacctacttacagaagacccccgccttatagggacccaagaccaccccccttccgacagggacggaaatggtggagaagcgacccctgcgggagaggcaccggacccctccccaatggcatctcgcctacgtgggagacgggag
ccccctgtggccgactccactacctcgcaggcattcccccctccgcgcaggaggaaacggacagcttcaatactggccgttctcctcttctgacctttacaactggaaaaataataacccttctttttctgaagatccaggtaaactgacagctct
gatcgagtctgtcctcatcacccatcagcccacctgggacgactgtcagcagctgttggggactctgctgaccggagaagaaaaacaacgggtgctcttagaggctagaaaggcggtgcggggcgatgatgggcgccccactcaactg
cccaatgaagtcgatgccgcttttcccctcgagcgcccagactgggattacaccacccaggcaggtaggaaccacctagtccactatcgccagttgctcctagcgggtctccaaaacgcgggcagaagccccaccaatttggccaaggt
aaaaggaataacacaagggcccaatgagtctccctcggccttcctagagagacttaaggaagcctatcgcaggtacactccttatgaccctgaggacccagggcaagaaactaatgtgtctatgtctttcatttggcagtctgccccagac
attgggagaaagttagagaggttagaagatttaaaaaacaagacgcttggagatttggttagagaggcagaaaagatctttaataaacgagaaaccccggaagaaagagaggaacgtatcaggagagaaacagaggaaaaagaa
gaacgccgtaggacagaggatgagcagaaagagaaagaaagagatcgtaggagacatagagagatgagcaagctattggccactgtcgttagtggacagaaacaggatagacagggaggagaacgaaggaggtcccaactc
gatcgcgaccagtgtgcctactgcaaagaaaaggggcactgggctaaagattgtcccaagaaaccacgaggacctcggggaccaagaccccagacctccctcctgaccctagatgactctggcggctcacttcaactgcctccacttg
aaagactgacactgggatcattacaattacctcctttagaacgattaacactcggttcactacagcttccgcctcttgagagattgacattaacgtccacgctgctaatggagaactcgtctggagattacaaagacgatgacgataagaaac
ggacagccgacggaagcgagttcgagtcaccaaagaagaagcggaaagtcGGaaacaattcccagggaagagtgaccttcgaggatgtcactgtgaacttcacccagggggagtggcagcggctgaatcccgaacagagaaa
cttgtacagggatgtgatgctggagaattacagcaaccttgtctctgtgggacaaggggaaaccaccaaacccgatgtgatcttgaggttggaacaaggaaaggagccatggttggaggaagaggaagtgctgggaagtggccgtgca
gaaaaaaatggggacattggagggcagatttggaagccaaaggatgtgaaagagagtctctacccaactttcttgtacaaagtggttggtggatctggaggttctcccaagaagaagcgcaaggtgggacgcgtctgcaggatatcaag
cttgcggtaccgcggcccgggatcgccaccatggacaagaagtacagcatcggcctggccatcggcaccaactctgtgggctgggccgtgatcaccgacgagtacaaggtgcccagcaagaaattcaaggtgctgggcaacaccg
accggcacagcatcaagaagaacctgatcggcgccctgctgttcgacagcggagaaacagccgaggccacccggctgaagagaaccgccagaagaagatacaccagacggaagaaccggatctgctatctgcaagagatcttc
agcaacgagatggccaaggtggacgacagcttcttccacagactggaagagtccttcctggtggaagaggataagaagcacgagcggcaccccatcttcggcaacatcgtggacgaggtggcctaccacgagaagtaccccaccat
ctaccacctgagaaagaaactggtggacagcaccgacaaggccgacctgcggctgatctatctggccctggcccacatgatcaagttccggggccacttcctgatcgagggcgacctgaaccccgacaacagcgacgtggacaagct
gttcatccagctggtgcagacctacaaccagctgttcgaggaaaaccccatcaacgccagcggcgtggacgccaaggccatcctgtctgccagactgagcaagagcagacggctggaaaatctgatcgcccagctgcccggcgaga
agaagaatggcctgttcggcaacctgattgccctgagcctgggcctgacccccaacttcaagagcaacttcgacctggccgaggatgccaaactgcagctgagcaaggacacctacgacgacgacctggacaacctgctggcccaga
tcggcgaccagtacgccgacctgtttctggccgccaagaacctgtccgacgccatcctgctgagcgacatcctgagagtgaacaccgagatcaccaaggcccccctgagcgcctctatgatcaagagatacgacgagcaccaccagg
acctgaccctgctgaaagctctcgtgcggcagcagctgcctgagaagtacaaagagattttcttcgaccagagcaagaacggctacgccggctacatcgatggcggagccagccaggaagagttctacaagttcatcaagcccatcctg
gaaaagatggacggcaccgaggaactgctcgtgaagctgaacagagaggacctgctgcggaagcagcggaccttcgacaacggcagcatcccccaccagatccacctgggagagctgcacgccattctgcggcggcaggaaga
ttttacccattcctgaaggacaaccgggaaaagatcgagaagatcctgaccttccgcatcccctactacgtgggccctctggccaggggaaacagcagattcgcctggatgaccagaaagagcgaggaaaccatcacccccctggaac
ttcgaggaagtggtggacaagggcgccagcgcccagagcttcatcgagcggatgaccaacttcgataagaacctgcccaacgagaaggtgctgcccaagcacagcctgctgtacgagtacttcaccgtgtacaacgagctgaccaa
agtgaaatacgtgaccgagggaatgagaaagcccgccttcctgagcggcgagcagaaaaaagccatcgtggacctgctgttcaagaccaaccggaaagtgaccgtgaagcagctgaaagaggactacttcaagaaaatcgagtgc
ttcgactccgtggaaatctccggcgtggaagatcggttcaacgcctccctgggcacataccacgatctgctgaaaattatcaaggacaaggacttcctggacaatgaggaaaacgaggacattctggaagatatcgtgctgaccctgaca
ctgtttgaggacagagagatgatcgaggaacggctgaaaacctatgcccacctgttcgacgacaaagtgatgaagcagctgaagcggcggagatacaccggctggggcaggctgagccggaagctgatcaacggcatccgggaca
agcagtccggcaagacaatcctggatttcctgaagtccgacggcttcgccaacagaaacttcatgcagctgatccacgacgacagcctgacctttaaagaggacatccagaaagcccaggtgtccggccagggcgatagcctgcacg
agcacattgccaatctggccggcagccccgccattaagaagggcatcctgcagacagtgaaggtggtggacgagctcgtgaaagtgatgggccggcacaagcccgagaacatcgtgatcgaaatggccagagagaaccagacca
cccagaagggacagaagaacagccgcgagagaatgaagcggatcgaagagggcatcaaagagctgggcagccagatcctgaaagaacaccccgtggaaaacacccagctgcagaacgagaagctgtacctgtactacctgc
agaatgggcgggatatgtacgtggaccaggaactggacatcaaccggctgtccgactacgatgtggacgctatcgtgcctcagagctttctgaaggacgactccatcgataacaaagtgctgactcggagcgacaagaaccggggca
agagcgacaacgtgccctccgaagaggtcgtgaagaagatgaagaactactggcgccagctgctgaatgccaagctgattacccagaggaagttcgacaatctgaccaaggccgagagaggcggcctgagcgaactggataagg
ccggcttcatcaagagacagctggtggaaacccggcagatcacaaagcacgtggcacagatcctggactcccggatgaacactaagtacgacgagaacgacaaactgatccgggaagtgaaagtgatcaccctgaagtccaagct
ggtgtccgatttccggaaggatttccagttttacaaagtgcgcgagatcaacaactaccaccacgcccacgacgcctacctgaacgccgtcgtgggaaccgccctgatcaaaaagtaccctaagctggaaagcgagttcgtgtacggcg
actacaaggtgtacgacgtgcggaagatgatcgccaagagcgagcaggaaatcggcaaggctaccgccaagtacttcttctacagcaacatcatgaactttttcaagaccgagattaccctggccaacggcgagatccggaagcggc
ctctgatcgagacaaacggcgaaacaggcgagatcgtgtgggataagggccgggactttgccaccgtgcggaaagtgctgtctatgccccaagtgaatatcgtgaaaaagaccgaggtgcagacaggcggcttcagcaaagagtcta
tcctgcccaagaggaacagcgacaagctgatcgccagaaagaaggactgggaccctaagaagtacggcggcttcgacagccccaccgtggcctattctgtgctggtggtggccaaagtggaaaagggcaagtccaagaaactgaa
gagtgtgaaagagctgctggggatcaccatcatggaaagaagcagcttcgagaagaatcccatcgactttctggaagccaagggctacaaagaagtgaaaaaggacctgatcatcaagctgcctaagtactccctgttcgagctggaa
aacggccggaagagaatgctggcctctgccggcgaactgcagaagggaaacgaactggccctgccctccaaatatgtgaacttcctgtacctggccagccactatgagaagctgaagggctcccccgaggataatgagcagaaaca
gctgtttgtggaacagcacaaacactacctggacgagatcatcgagcagatcagcgagttctccaagagagtgatcctggccgacgctaatctggacaaggtgctgagcgcctacaacaagcacagagacaagcctatcagagagca
ggccgagaatatcatccacctgtttaccctgaccaatctgggagcccctgccgccttcaagtactttgacaccaccatcgaccggaagaggtacaccagcaccaaagaggtgctggacgccaccctgatccaccagagcatcaccggc
ctgtacgagacacggatcgacctgtctcagctgggaggcgactctggcggctcaaaaagaaccgccgacggcagcgaattcgagtctcccaagaagaagaggaaagtcggctctggccctgccgctaagagagtgaagctggactg
agatctttttccctctgccaaaaattatggggacatcatgaagccccttgagcatctgacttctggctaataaaggaaatttattttcattgcaatagtgtgttggaattttttgtgtctctca

**Supplementary Table 2. Target (with PAM) and primer sequences.**

| Name | Sequence 5' – 3' | Target spacer (with PAM) | Species |
|---|---|---|---|
| *RNF2* NGS Forward | ACACTCTTTCCCTACACGACGCTCTTCCGATCTATTTCCAGCAATGTCTCAGG | GTCATCTTAGTCATTACCTGAGG | Human |
| *RNF2* NGS Reverse | GTGACTGGAGTTCAGACGTGTGCTCTTCCGATCTGCCAACATACAGAAGTCAGGAA | GTCATCTTAGTCATTACCTGAGG | Human |
| *EMX1* NGS Forward | ACACTCTTTCCCTACACGACGCTCTTCCGATCTGGACAAAGTACAAACGGCAGA | GAGTCCGAGCAGAAGAAGAAGGG | Human |
| *EMX1* NGS Reverse | GTGACTGGAGTTCAGACGTGTGCTCTTCCGATCTAGTGGCCAGAGTCCAGCTT | GAGTCCGAGCAGAAGAAGAAGGG | Human |
| *CD244* NGS Forward | ACACTCTTTCCCTACACGACGCTCTTCCGATCTACATCCAGCTGCCCACAT | GACCATGTGGTTAGCATCTCGGG | Human |
| *CD244* NGS Reverse | GTGACTGGAGTTCAGACGTGTGCTCTTCCGATCTGGAAGGCAAAGAGCCATTCT | GACCATGTGGTTAGCATCTCGGG | Human |
| *HEK2* NGS Forward | ACACTCTTTCCCTACACGACGCTCTTCCGATCTGGAATGAATGGATTCCTTGG | GAACACAAAGCATAGACTGCGGG | Human |
| *HEK2* NGS Reverse | GTGACTGGAGTTCAGACGTGTGCTCTTCCGATCTTAGTTAAGAACACGTTTAAA | GAACACAAAGCATAGACTGCGGG | Human |
| *AAVS1* NGS Forward | ACACTCTTTCCCTACACGACGCTCTTCCGATCTAGGATCCTCTCTGGCTCCAT | CCCTAGTGGCCCCACTGTGGGG | Human |
| *AAVS1* NGS Reverse | GTGACTGGAGTTCAGACGTGTGCTCTTCCGATCTCGGTTAATGTGGCTCTGGTT | CCCTAGTGGCCCCACTGTGGGG | Human |
| *HEK3* NGS Forward | ACACTCTTTCCCTACACGACGCTCTTCCGATCTAAACGCCCATGCAATTAGTC | GGCCCAGACTGAGCACGTGATGG | Human |
| *HEK3* NGS Reverse | GTGACTGGAGTTCAGACGTGTGCTCTTCCGATCTCCAGCCAAACTTGTCAACC | GGCCCAGACTGAGCACGTGATGG | Human |
| *HBG* NGS Forward | ACACTCTTTCCCTACACGACGCTCTTCCGATCTCGGCTGACAAAAGAAGTCCT | GCATTGAGATAGTGTGGGGAAGG | Human |
| *HBG* NGS Reverse | GTGACTGGAGTTCAGACGTGTGCTCTTCCGATCTAGGCTATTGGTCAAGGCAAG | GCATTGAGATAGTGTGGGGAAGG | Human |
| *MLH1* NGS Forward | ACACTCTTTCCCTACACGACGCTCTTCCGATCTGCTGAAGGAAGAACGTGAGC | TCGGCGGCTGGACGAGACAGTGG | Human |
| *MLH1* NGS Reverse | GTGACTGGAGTTCAGACGTGTGCTCTTCCGATCTCGACTCCCTCCGTACCAGT | TCGGCGGCTGGACGAGACAGTGG | Human |
| *HEK2* OT1 NGS Forward | **ACACTCTTTCCCTACACGACGCTCTTCCGATCTGTGTGGAGAGTGAGTAAGCCA** | GAACACAATGCATAGATTGCCGG | Human |
| *HEK2* OT1 NGS Reverse | **GTGACTGGAGTTCAGACGTGTGCTCTTCCGATCTACGGTAGGATGATTTCAGGCA** | GAACACAATGCATAGATTGCCGG | Human |
| *HEK3* OT1 NGS Forward | **ACACTCTTTCCCTACACGACGCTCTTCCGATCTTCCCCTGTTGACCTGGAGAA** | CACCCAGACTGAGCACGTGCTGG | Human |
| *HEK3* OT1 NGS Reverse | **GTGACTGGAGTTCAGACGTGTGCTCTTCCGATCTCACTGTACTTGCCCTGACCA** | CACCCAGACTGAGCACGTGCTGG | Human |
| *HEK3* OT2 NGS Forward | **ACACTCTTTCCCTACACGACGCTCTTCCGATCTTTGGTGTTGACAGGGAGCAA** | GACACAGACTGGGCACGTGAGGG | Human |
| *HEK3* OT2 NGS Reverse | **GTGACTGGAGTTCAGACGTGTGCTCTTCCGATCTCTGAGATGTGGGCAGAAGGG** | GACACAGACTGGGCACGTGAGGG | Human |
| *HEK3* OT3 NGS Forward | **ACACTCTTTCCCTACACGACGCTCTTCCGATCTTGAGAGGGAACAGAAGGGCT** | AGCTCAGACTGAGCAAGTGAGGG | Human |
| *HEK3* OT3 NGS Reverse | **GTGACTGGAGTTCAGACGTGTGCTCTTCCGATCTGTCCAAAGGCCCAAGAACCT** | AGCTCAGACTGAGCAAGTGAGGG | Human |

| Name | Sequence 5' – 3' | Target spacer (with PAM) | Species |
|---|---|---|---|
| *Vegfa* NGS Forward | ACACTCTTTCCCTACACGACGCTCTTCCGATCTCAAATCTGGGTGGCGATAGA | CTCCTGGAAGATGTCCACCAGGG | Mouse |
| *Vegfa* NGS Reverse | GTGACTGGAGTTCAGACGTGTGCTCTTCCGATCTCTCCAGGGCTTCATCGTTACA | CTCCTGGAAGATGTCCACCAGGG | Mouse |
| *Rosa*26 NGS Forward | ACACTCTTTCCCTACACGACGCTCTTCCGATCTTCCCAAAGTCGCTCTGAGTT | GGCGGTCCTCAGAAGCCAGGAGG | Mouse |
| *Rosa*26 NGS Reverse | GTGACTGGAGTTCAGACGTGTGCTCTTCCGATCTCTTTAAGCCTGCCCAGAAGA | GGCGGTCCTCAGAAGCCAGGAGG | Mouse |
| *Dnmt*1 NGS Forward | ACACTCTTTCCCTACACGACGCTCTTCCGATCTCCTTCGGGCATAGCATGGT | CGGGCTGGAGCTGTTCGCGCTGG | Mouse |
| *Dnmt*1 NGS Reverse | GTGACTGGAGTTCAGACGTGTGCTCTTCCGATCTATATATGCCTCGGCATCGGT | CGGGCTGGAGCTGTTCGCGCTGG | Mouse |
| *Rpe*65 NGS Forward | ACACTCTTTCCCTACACGACGCTCTTCCGATCTCCACCAAATTGCATAAGAAAAA | ACTGGCAGTCTCCTCTGATGTGG | Mouse |
| *Rpe*65 NGS Reverse | GTGACTGGAGTTCAGACGTGTGCTCTTCCGATCTCCTTCTGTGGTATGTGACAT | ACTGGCAGTCTCCTCTGATGTGG | Mouse |
| *Ar* NGS Forward | ACACTCTTTCCCTACACGACGCTCTTCCGATCTGGACCATGTTTTACCCATCG | TCTCACTTGTGGCAGCTGCAAGG | Mouse |
| *Ar* NGS Reverse | GTGACTGGAGTTCAGACGTGTGCTCTTCCGATCTTATCAGAATGCCCTGGAAGG | TCTCACTTGTGGCAGCTGCAAGG | Mouse |

| Name | Sequence 5' – 3' | Species |
|---|---|---|
| *HBG* qRT-PCR Forward | GCTGAGTGAACTGCACTGTGA | Human |
| *HBG* qRT-PCR Reverse | GAATTCTTTGCCGAAATGGA | Human |
| *MLH*1 qRT-PCR Forward | GTTCTCCGGGAGATGTTGCATA | Human |
| *MLH*1 qRT-PCR Reverse | TGGTGGTGTTGAGAAGGTATAACTTTG | Human |
